# Iterative model-based density improvement yields better atomic structures from cryo-EM maps

**DOI:** 10.1101/121913

**Authors:** Arjen J. Jakobi, Matthias Wilmanns, Carsten Sachse

## Abstract

Atomic models based on high-resolution density maps are the ultimate result of the cryo-EM structure determination process. Current cryo-EM model refinement procedures work with the experimental density map that remains constant throughout the process. Here, we introduce a general procedure that iteratively improves cryo-EM density maps based on prior knowledge of an atomic reference structure. The procedure optimizes contrast of cryo-EM densities by local amplitude scaling (LocScale) based on an atomic model without introducing model bias. We alternate the procedure with consecutive rounds of model refinement and tested it on four cryo-EM structures of TRPV1, β-galactosidase, γ-secretase and RNA polymerase III. We demonstrate that LocScale density improvement reveals previously undiscovered map features and improves the quality of atomic models. The presented approach enhances the interpretability of cryo-EM density maps and provides an implementation reminiscent of iterative density improvement as it is routinely employed in the refinement of X-ray crystallographic models.

## Introduction

Electron cryo-microscopy (cryo-EM) has been used as a method to visualize biological macromolecules in their native-hydrated state for more than three decades (Adrian et al., 1984). Major improvements in detector technology (Faruqi and Henderson, 2007; McMullan et al., 2016) and associated computational procedures recently transformed single-particle cryo-EM that since has been producing a plenitude of near-atomic resolution structures from specimens of lower symmetry (Allegretti et al., 2014; Amunts et al., 2014; Bai et al., 2013) and lower molecular weight than previously deemed possible (Bai et al., 2015; Liao et al., 2013; Merk et al., 2016). At this resolution, the reconstructed EM density maps contain sufficient detail to interpret the structure using atomic models. Atomic models are frequently used by biologists without expert structural expertise to interrogate function based on mechanistic hypotheses inferred from the structure. The building and refinement of atomic models thus present a critical step in the structure determination process and their accuracy depends on the quality of the EM density.

Sophisticated protocols for map interpretation as well as parameterization and validation of coordinate refinement in X-ray crystallography exist and these procedures have been adapted to work with cryo-EM maps (Amunts et al., 2014; Brown et al., 2015; Fromm et al., 2015; Hoffmann et al., 2015; Z. Wang et al., 2014). Coordinate refinement in X-ray crystallography typically depends on initial phase estimates that are iteratively improved (Agarwal and Isaacs, 1977; Grosse-Kunstleve et al., 2002; Lunin et al., 2002; Read, 1986; Wilson, 1942). As a result, crystallographic refinement successively improves map interpretability throughout the refinement procedure. By contrast, the cryo-EM experiment yields both amplitudes and phases directly from images and experimentally determined EM density can therefore be used as a constant minimization target (Grigorieff et al., 1996; Unwin, 2005; Yonekura et al., 2003).

Regardless of implementation, all model building and refinement procedures are guided by the structural information content present in the data, which is primarily defined by resolution. The commonly accepted procedure for resolution estimation in cryo-EM is the Fourier shell correlation (FSC), which measures the correlation of Fourier coefficients in resolution shells between independent map reconstructions (Harauz and van Heel, 1986; Saxton and Baumeister, 1982; van Heel et al., 1982). The reported map resolution corresponds to a pre-defined cut-off of the FSC curve (Rosenthal and Henderson, 2003). Currently, it is common practice to low-pass filter cryo-EM volumes at this resolution cut-off. Many of the recently determined EM density maps, however, contain regions that differ substantially from this overall resolution due to varying flexibility and/or subunit occupancy in different parts of the structure (Bai et al., 2013; Hoffmann et al., 2015; Zhao et al., 2015). In addition to inherent flexibility of macromolecular complexes, the quality of the reconstruction is poorer at the particle periphery when compared with the particle center due to limitations of current three-dimensional (3D) reconstruction algorithms. In providing an estimate of overall map resolution, the average FSC measurement falls short of describing such local resolution differences in different parts of the structure (Cardone et al., 2013).

In addition to resolution, density contrast is critical for map interpretation. High-resolution contrast in cryo-EM maps is attenuated by a resolution-dependent amplitude fall-off. Compensation is achieved by sharpening with a uniform map B-factor, which partially restores contrast loss in cryo-EM maps (Rosenthal and Henderson, 2003) and in low-resolution electron density maps from X-ray crystallography (DeLaBarre and Brunger, 2003; Jacobson and Wunderlich, 1961; Stehle and Harrison, 1996). This B-factor can be estimated numerically (DeLaBarre and Brunger, 2003; Rosenthal and Henderson, 2003) but in practice often needs to be fine-tuned empirically in an effort to generate optimal contrast. Alternative scaling procedures use the X-ray solution scattering (SAXS) curve of the specimen (Gabashvili et al., 2000; Schmid et al., 1999) or the radially averaged amplitude profile of the crystal structure (Falke et al., 2005) to correct the amplitudes accordingly. The desired outcome of all procedures is to enhance contrast such that it reveals expected molecular features at the given resolution while avoiding undue amplification of background noise. Depending on the resolution, such features comprise clearly defined signatures of secondary structure and amino acid side chains. With the wealth of recent near-atomic resolution cryo-EM reconstructions (Cheng, 2015; Kühlbrandt, 2014; Subramaniam et al., 2016), building and refinement of atomic models have become the ultimate steps of cryo-EM structure determination. Therefore, new approaches are required to optimally guide the process of sharpening based on prior knowledge of atomic models and to better integrate model refinement with map interpretation.

Crystallographic model refinement routinely uses prior information from atomic models to enhance the clarity of electron density maps. As the model improves, better phase estimates iteratively make the electron density more interpretable and ultimately provides more accurate structures. Current cryo-EM density interpretation does not make use of model information and model refinement strategies leave the target density unchanged. We here present a general procedure that utilizes prior model information to iteratively compute map representations better suited for interpretation and atomic model refinement than those obtained from existing approaches. The procedure optimizes map contrast by local amplitude scaling against an atomic structure and thereby implicitly accounts for differences in local resolution without introducing model bias. Based on several test cases from the EMDB model challenge (http://challenges.emdatabank.org), we show that model-based density improvement yields refined atomic models of better fit and geometry.

## Results

### Integrating cryo-EM density improvement with iterative atomic model refinement

We propose the following procedure to integrate cryo-EM density improvement with atomic model refinement: First, an initial atomic model is generated and atomic coordinates and B-factors are subsequently refined using existing procedures (Hoffmann et al., 2015; Murshudov, 2016; Z. Wang et al., 2014). In a second step, we use the refined coordinate model to generate a reference map used for local amplitude scaling (LocScale) of the cryo-EM density. The LocScale density contains enhanced local contrast and can be used as an improved target map for additional rounds of model building and refinement. The procedure can be iterated until convergence (**Figure 1**).

**Figure 1.**
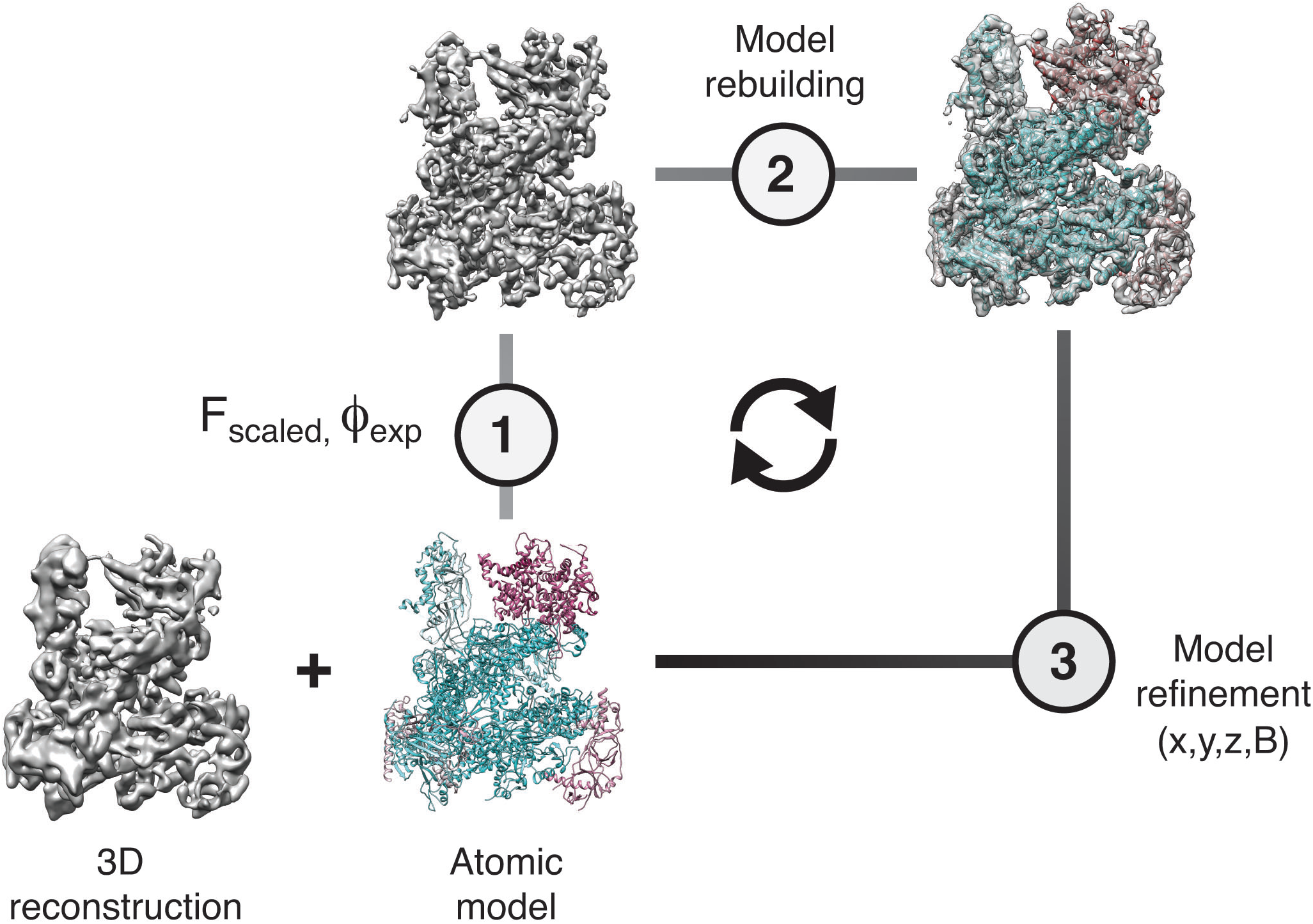
Cyclical workflow of cryo-EM density improvement using reference-based scaling alternating with model refinement. An atomic model is used as a reference to improve contrast in the original cryo-EM reconstruction employing local amplitude scaling (1). The resulting LocScale map contains enhanced structural detail and thus presents a better density for model building (2) and improved target for coordinate refinement (3). Scaling, building and refinement steps can be iterated to produce successively improved density maps and atomic models.

### Improved image contrast by local reference-based amplitude scaling

The central concept of the proposed method is the improvement of map contrast by local amplitude scaling. The appearance of map features does not only depend on the existence of amplitude and phase signal at the corresponding resolution, but also on the relative magnitude of these frequency components. This is particularly true for cryo-EM images that are affected by amplitude modulations due to the contrast-transfer function (Erickson and Klug, 1970). To examine the effect of various amplitude-based contrast enhancing procedures, we manipulated the amplitudes of a perfect two-dimensional (2D) test image of 512 × 512 pixel dimensions (Figure 2A). For illustration purposes, we substituted the amplitudes of the image with random Gaussian noise and computed the inverse Fourier transform of the hybrid image. Although this image still has perfect phases and the overall image information is largely conserved, it shows poor contrast and thus makes feature discrimination difficult, and in part impossible (Figure 2B). As reflected in the radial amplitude profile of the original image, image amplitudes decay exponentially towards higher frequencies. Amplitudes of cryo-EM maps decay much faster due to imaging imperfections (Henderson, 1992). Scaling amplitudes by an approximated exponential fall-off restores the overall magnitude distribution at low and high frequencies and improves image contrast (Figure 2C). This procedure is equivalent to the widely used cryo-EM map sharpening procedure with a numerically estimated B-factor. Better contrast can be obtained by more accurately restoring amplitude differences between adjacent frequency components. This can be achieved by scaling amplitudes against the radial average of the original image, equivalent to so-called reference-based scaling of cryo-EM maps using an X-ray model or solution scattering profile (Figure 2D). Moreover, as the radially averaged amplitude profile sums many features across the image, it is evident that the relative distribution of adjacent frequency components may not be optimally restored by global amplitude correction. We therefore tested whether local image scaling can yield more accurate amplitude distributions. To this end, we implemented a tile-based scaling procedure, in which the amplitudes of 128 × 128 pixel rolling windows were scaled against the local radial average of the respective reference window and the central pixels of the scaled windows make up the resulting image. In this image, high-resolution contrast including directional amplitude components is better restored when compared with that scaled against the global average, and the power spectrum more closely resembles that of the original image (Figure 2E). This 2D example highlights the importance of relative amplitude scaling for local image contrast and shows that contrast restoration by tile-based amplitude scaling against a local reference should be preferred over compensating amplitude decay by an exponential or an overall fall-off approximation.

**Figure 2.**
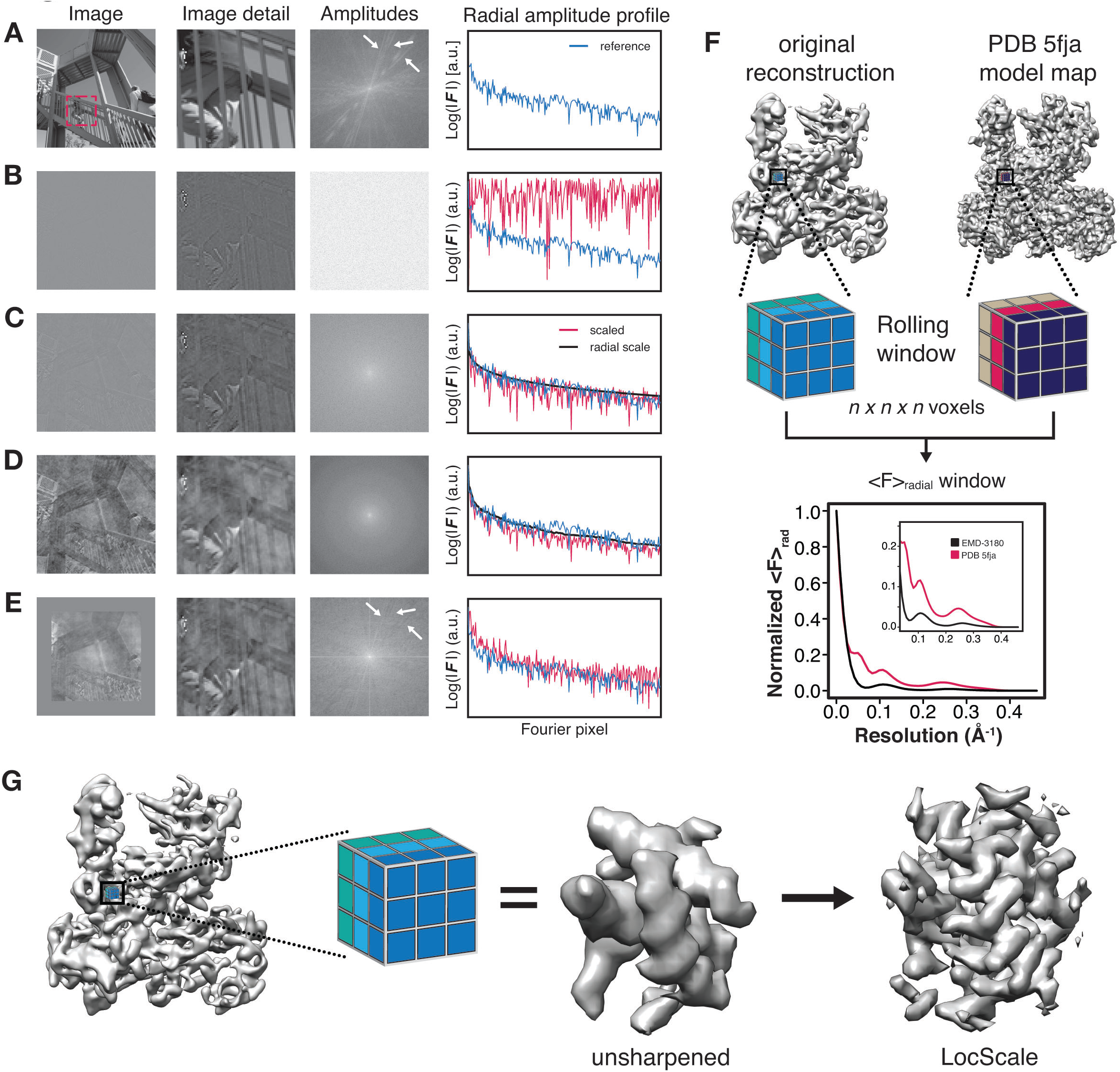
Illustration of local amplitude scaling (LocScale) using a 2D test image and generation of 3D LocScale density maps. (**A**) The original image (left), a magnified image detail (center left), the overall amplitude spectrum (center right) and the radially averaged amplitude profile (right) are displayed in panels along columns, respectively. Applied procedures are shown along rows. The radial amplitude profile of the reference, the scaled image and the applied scale are shown in blue, red and black, respectively. (**B**) Hybrid image resulting when amplitudes were replaced by random Gaussian noise and combined with phases from the original image in (B). The fall-off of the amplitude profile of the hybrid image (red) is flat over the entire frequency range and hampers visual perception of image contrast. (**C**) Hybrid image from (C) when random amplitudes were scaled by a negative exponential estimated from the amplitude fall-off of the original image. (**D**) Hybrid image from (C) with random amplitudes scaled by the average radial fall-off profile from (B). (**E**) Hybrid image from (C) with amplitudes scaled locally to reference amplitudes from (B) computed in tiles of 128x128 pixel rolling windows. Note that diagonal lines in the amplitude spectrum (white arrows) arising from periodic image features are optimally recovered. Compare with the power spectrum of the original image. (**F**) Schematic illustration of the LocScale procedure for a Pol III density map. Two equivalent map segments (rolling windows) are extracted from the original 3D reconstruction of EMD-3180 (top) and a map simulated from the atomic model (PDB ID 5fja) (bottom). For each rolling window, the radially averaged structure factor profile is computed and used to scale the amplitudes of the corresponding window of the original reconstruction. Note, the fine structure of the model amplitudes has a characteristic profile in the resolution range <10Å due to protein secondary structure and deviates from a simple exponential fall-off (inset). (**G**)Effect of amplitude scaling illustrated for an exemplary density window. The density contained within a window cube is shown before (unsharpened) and after (LocScale) application of amplitude scaling. The central voxel of the rolling window is assigned the map value after amplitude scaling. The procedure is repeated by moving the window along the map until each voxel has been assigned a density value based on the locally estimated contrast.

### Local amplitude scaling optimizes contrast in cryo-EM maps

Due to the contrast benefit of local amplitude scaling in photographic images, we set out to assess whether this strategy improves contrast restoration in cryo-EM density maps. Contrary to the 2D image with uniform resolution, cryo-EM maps display local resolution variations in addition to the steep amplitude decay towards high resolution. As a consequence, it is conceivable that cryo-EM maps require locally adjusted sharpening levels to avoid over-amplification of noise and dampening of high-resolution features. We therefore implemented a procedure to perform local map sharpening for cryo-EM reconstructions analogous to the tile-based approach for the 2D test image outlined above. In this procedure, we locally scale the experimental amplitudes to match the radially averaged amplitude profile of a map generated from a refined atomic reference model (Figure 2F). Within a rolling 3D density window, the procedure adapts the relative amplitude levels required to optimize local contrast, by imposing the expected amplitude profile for the structure contained within it and by acting as a convoluted Gaussian low-pass filter that reduces background noise (Figure 2G). The first effect is related to the amplitude fall-off of atoms organized in secondary structure and other periodic elements inherent to polypeptide chains and nucleic acids (Figure 2–figure supplement 1 A-C). This amplitude fall-off is markedly different to the exponential decay of randomly distributed atoms, which is approximated and compensated for by common sharpening procedures. The second effect of low-pass filtration arises from the local distribution of atomic B-factors that determines the steepness of the amplitude fall-off with increasing resolution (Figure 2–figure supplement 1 D/E). Performing the procedure across the density map results in a single map with optimized contrast. Throughout the manuscript, we refer to these maps as LocScale maps.

### LocScale maps improve interpretability of cryo-EM maps

We illustrate the results of the LocScale procedure for cryo-EM maps of Pol III (EMD-3180) and TRPV1 channel (EMD-5778). Overall, the LocScale map of Pol III appears cleaner than the uniformly sharpened map and secondary structure elements are readily recognizable (Figure 3A, left). At first sight, the Pol III LocScale map may appear less featured at the peripheral subunits when compared with EMD-3180. However, closer inspection reveals that features in the deposited map in fact are sharpening artifacts inconsistent with molecular structure (Figure 3A, upper panel). By contrast, LocScale density improvement eliminates such artifacts resulting from incorrect amplitude scaling (Figure 3A, lower panel). In a second example, we compared the LocScale map of the TRPV1 channel with the deposited EMD-5778 map. This comparison reveals more clearly resolved α-helical pitch and well-defined side chain densities in the LocScale map (Figure 3B, lower panel), which are not as readily discernible in the uniformly sharpened and filtered map (Figure 3B, upper panel). In effect, the optimized contrast enhances high-resolution features from map regions at higher than average resolution in addition to dampening noise in regions at lower than average resolution. LocScale density improvement enhances interpretability of cryo-EM density maps by restoring optimal local map contrast.

**Figure 3.**
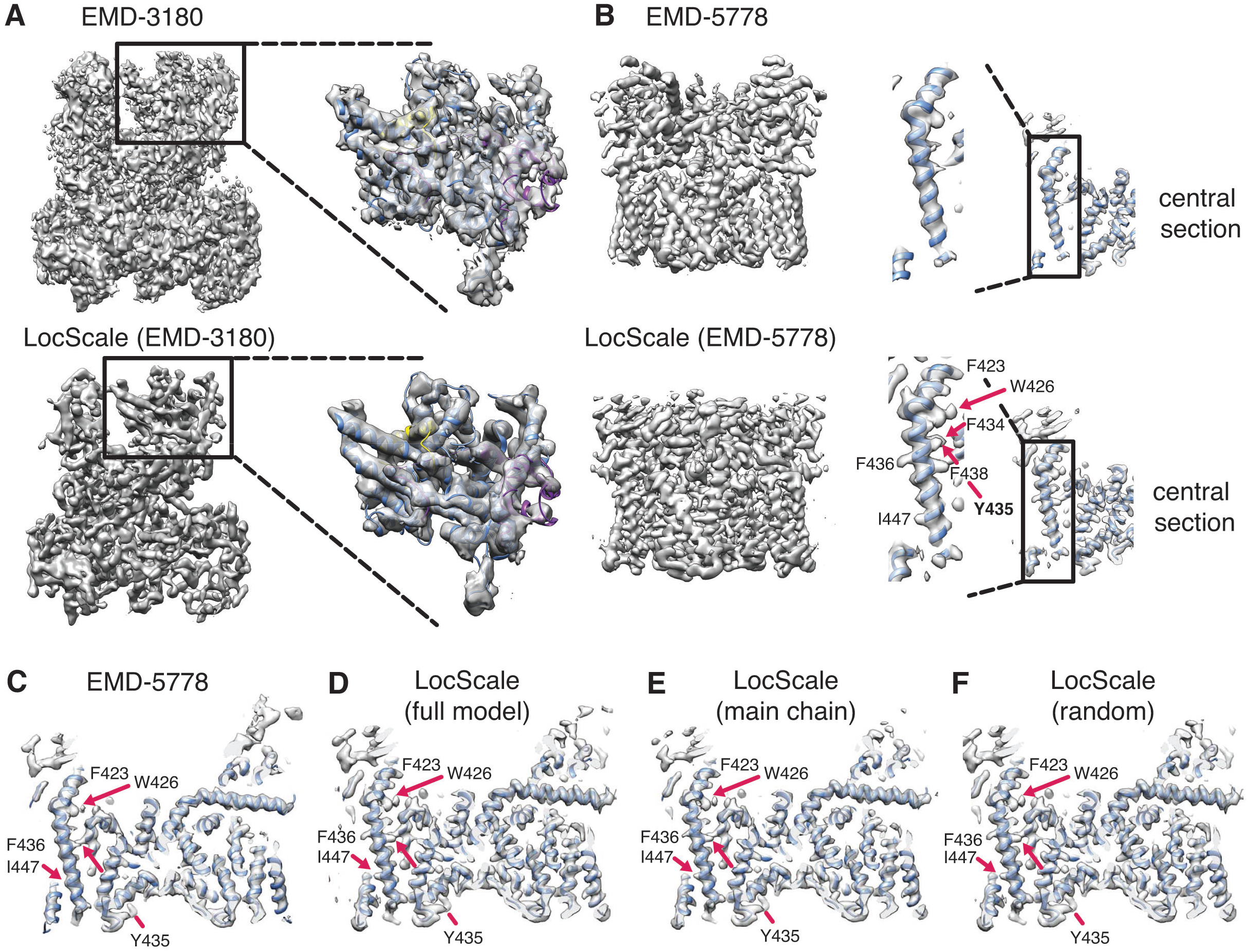
Local amplitude scaling (LocScale) optimizes cryo-EM map contrast and does not introduce model bias. **(A)** and **(B)** Illustration of the LocScale density improvement for initially over-sharpened (**A**, RNA Pol III; EMD-3180) and under-sharpened maps (**B**, TRPV1; EMD-5778). The maps after application of global (Guinier) sharpening (top) are shown in comparison with its LocScale equivalent (bottom). The LocScale map contains recognizable secondary structure. Zoomed-in inset: (A) The globally sharpened map shows undue noise amplification rather than true molecular features of the peripheral subunits as visualized in the LocScale map. Central section: (B) When global sharpening levels were underestimated, LocScale map reveals previously invisible features such as defined side chain densities (arrows). **(C)** Central slice through deposited TRPV1 map (EMD-5778) with superimposed atomic model in ribbon representation. **(D)** LocScale map obtained with the all-atom reference model shows improved definition of helical pitch and side-chain densities become visible. **(E)** LocScale map obtained using a reference model limited to main chain atoms reveals side chain densities even though not present in the reference. **(F)** LocScale map obtained using a random reference amplitude profile at comparable resolution. The resulting map features are similarly well recovered when compared with those obtained by the original procedure in (B).

### Atomic model does not introduce reference bias on LocScale density

To ascertain that these features originate from enhanced contrast and not from model bias, we used the TRPV1 structure to compute multiple LocScale maps using different reference amplitudes, i.e. we tested the effect of the reference model on the density modification procedure (Figure 3C-F). In a first experiment, we performed local amplitude scaling using a reference map that was computed from a main chain-only model (Figure 3E). In a second experiment, we used randomized reference windows with approximately equivalent resolution that originated from completely different regions of the reference map (Figure 3F). The resulting LocScale maps in both cases consistently showed visible side chain densities in the well-ordered transmembrane region, confirming that these features indeed are the result of enhanced contrast and not of bias from the reference model. The LocScale procedure only adapts amplitudes by scaling according to a local radial average computed from the reference structure. As it ignores model phases and the scaled amplitudes are recombined with the experimentally determined map phases, the method does not introduce bias towards the model.

In addition to model coordinate bias, we tested the effect of atomic B-factors associated with the reference coordinates on the resulting LocScale map. As the atomic B-factor distribution determines the steepness of the reference amplitude fall-off within the scaling window, we reasoned that correctly estimated atomic B-factors would be important for optimal performance of the LocScale density improvement. If refinement proceeds robustly, atomic B-factors should be largely in correspondence with local resolution estimates on a per residue basis, as exemplified for Pol III (EMD-3180 and PDB ID 5fja; Figure 3–figure supplement 1A/B). To examine the effect of estimated B-factors on the scaling procedure, we generated LocScale maps of EMD-3180 based on reference maps from models whose atoms were assigned a series of uniform B-factors ranging from 0 to 300 Å^2^ (Figure 3–figure supplement 1C). Where atomic B-factors were underestimated (B ≤ 50 Å^2^) for model subunits located in the low-resolution map region at the particle periphery, the shallow fall-off in the scaling reference led to density fragmentation in the scaled map (Figure 3–figure supplement 1D-F). Conversely, a local amplitude profile that falls off too steeply by overestimated atomic B-factors within the core of the molecule (B ≥ 200 Å^2^) resulted in blurring of well resolved secondary structure elements in the respective scaled map (Figure 3–figure supplement 1D/E). Therefore, using incorrectly estimated atomic B-factors for LocScale can lead to a suboptimal representation of density features, but does not introduce bias from the reference. For correctly estimated atomic B-factor distributions, the level of blurring and sharpening is locally optimized for each region leading to optimal density contrast in the LocScale map (Figure 3–figure supplement 1C-F).

### LocScale maps facilitate model building and reveal novel structural detail

To test whether LocScale maps can aid model building and refinement we chose three cryo-EM structures from the 2015/2016 EMDB model challenge (http://challenges.emdatabank.org), TRPV1 channel (EMD-5778), γ-secretase (EMD-3061) and β-galactosidase (EMD-2984), and one additional test case from our own lab, Pol III (EMD-3180). All four systems pose unique challenges: Pol III displays considerable variation in local resolution across the map ranging from good (<4Å) to very low resolution (>8Å; see above). γ-secretase (Bai et al., 2015) contains large local resolution differences between the protein core and embedding amphipols. The TRPV1 channel (Liao et al., 2013) displays substantial resolution differences between the transmembrane and extracellular regions. β-galactosidase (Bartesaghi et al., 2015) is an example of a structure resolved at an exceptionally high resolution (2.2 Å). These examples allowed us to test the effectiveness of the procedure for a resolution range from ∼8 to 2 Å. To start with comparable reference models, we first generated uniformly filtered and sharpened maps and performed one round of atomic B-factor refinement using the deposited PDB coordinates as starting models. For further analysis, we computed LocScale maps using the resulting reference models as described above.

We first assessed the validity of our procedure by comparing the deposited map of β-galactosidase (EMD-2984) with the map obtained by the LocScale procedure. The density of EMD-2984 is of such exceptional quality that individual water molecules can be placed, and characteristic doughnut-shaped densities of aromatic rings are revealed in the best-resolved regions of the map (Bartesaghi et al., 2015). Visualizing these high-resolution features in the deposited map required sharpening levels that result in a density with high overall noise levels (Figure 4A). Qualitatively, the LocScale map appears less noisy when compared with the deposited map, which is supported by the smooth fall-off of the radially averaged power spectrum profile (Figure 4–figure supplement 1A). At the same time, the LocScale map does preserve the reported high-resolution details such as ordered water molecules and characteristic density holes of aromatic rings (Figure 4A-B). While most of the structure shows an excellent fit between density and the deposited coordinate model (PDB ID 5a1a), we noted that one particular region (V728–H735; Figure 4C inset) displayed poor real-space correlation and the model in this region contains unusually high B-factors relative to other parts of the structure. Closer inspection of the deposited model and map revealed that density in this region is fragmented and precludes confident modelling of the backbone conformation (Figure 4D). By contrast, the LocScale density showed a clear backbone trace and reveals that the deposited coordinates in this region are incompatible with the density (Figure 4E). The LocScale map allowed us to rebuilt this residue stretch to better satisfy the density (Figure 4F, Figure 4–figure supplement 1B). To verify our rebuilt conformation, we compared both PDB ID 5a1a and the rebuilt model with the 1.6 Å resolution crystal structure of *E.coli* β-galactosidase ((Wheatley et al., 2015), PDB ID 4ttg). The V728–H735 conformation built and refined using the LocScale map is in closer agreement with the high-resolution crystal structure than the deposited model (Figure 4–figure supplement 1C/D) and converges after three LocScale iterations alternating with model refinement (Figure 4–figure supplement 1E). In EMD-2984, the fragmented density and high noise level in this region appears to have misguided model building.

**Figure 4.**
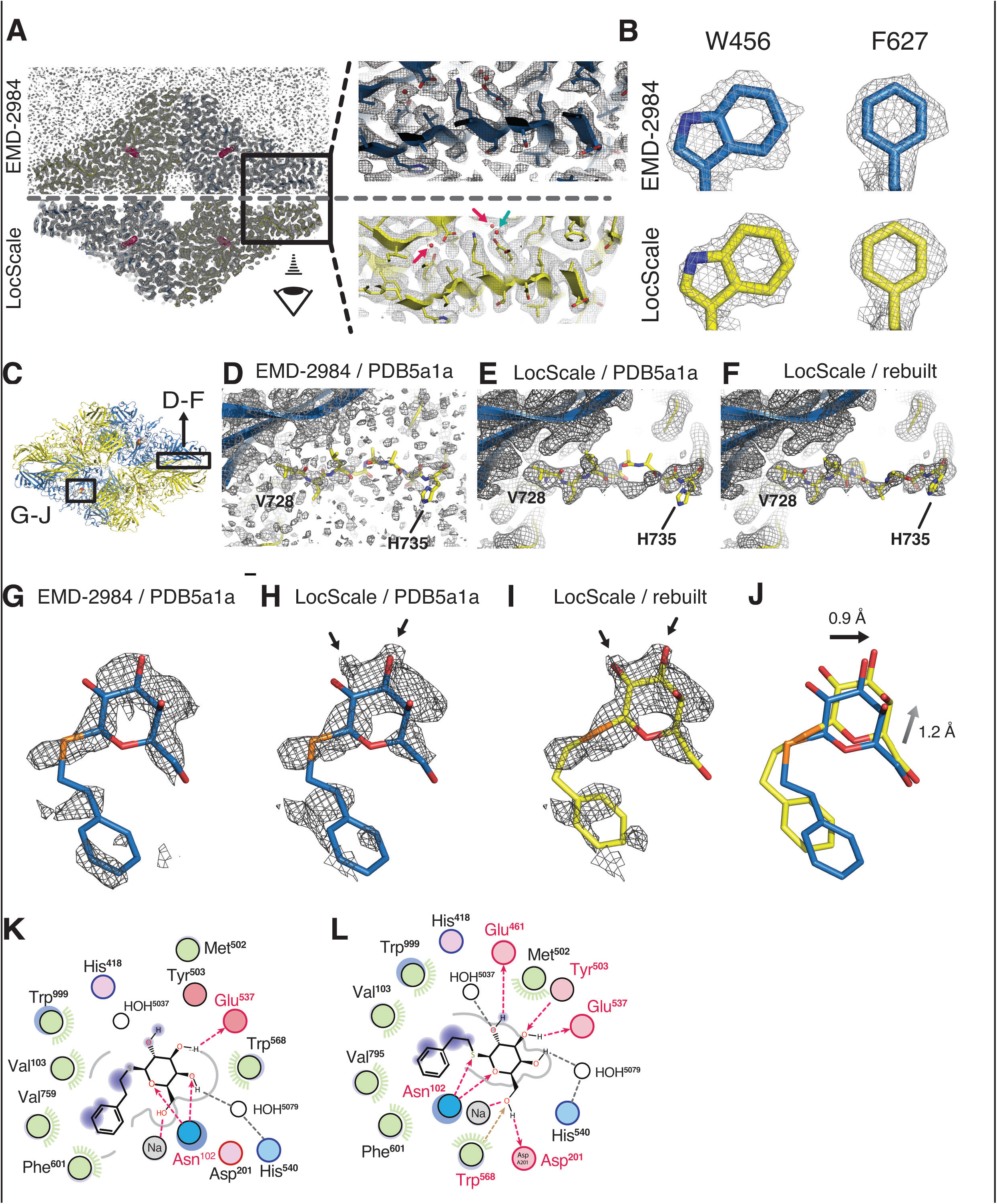
Application of the LocScale procedure to the 2.2 Å resolution cryo-EM structure of β-galactosidase. **(A)** Overall comparison of uniformly sharpened (EMD-2984) (top) and LocScale map (bottom) for the β-galactosidase structure. EMD-2984 displays visible noise at the periphery of the molecule and its surrounding solvent, whereas the LocScale map contains reduced noise levels in corresponding regions. Zoomed-in regions in the protein core showing modeled water molecules for which density is retained in the LocScale map (arrows). **(B)** High-resolution details such as doughnut-shaped densities for aromatic side chains W456 and F627 in the EMD-2984 map (top) are consistently retained in the LocScale map (bottom). Maps are shown at equivalent isosurface levels displayed at a 1.6 Å carve radius. **(C)** Cartoon representation of the β-galactosidase tetramer (PDB ID 5a1a) colored by symmetry-related subunits (yellow and blue). **(D-F)** Density around the peripheral loop region comprising residues V728–H735 (see (C) for overall location) fragmented in the deposited map, hampering unambiguous tracing of the backbone conformation. Deposited conformation in PDB ID 5a1a superimposed on the EMD-2984 map (D) and LocScale map (E). Note reduced noise levels and continuous density providing an unambiguous backbone trace in the LocScale map. The LocScale density is not compatible with the modelled conformation of PDB ID 5a1a, but supports an alternative conformation of V728–H735 shown superimposed on the LocScale map (F). (**G-J**) Density for the PETG ligand bound in the active site as observed in EMD-2984 (G) and LocScale map (H) superposed on the PETG conformation from PDB ID 5fja shown in stick representation. Enhanced density features of the hydroxyl substituents in the hexapyranosyl ring become visible in the LocScale map (arrows) and require re-arrangement of the ligand orientation in the active site **(I and J)**. **(K and L)** FLEV ligand environment computed from PDB ID (5fja) and the model after refinement against the LocScale map. Re-orientation of the ligand based on the LocScale map increases hydrogen-bond interactions in the active site. Note also the significantly decreased substitution contour (gray line) for the LocScale conformation, indicating tighter apposition in the active site.

The β-galactosidase cryo-EM structure contains the inhibitor phenylethyl-β-D-thiogalactopyranoside (PETG) in the active site. When analyzing the ligand density in the active site binding pocket, we realized that the LocScale map showed additional density features suggestive of hydroxyl substituents at the hexa-pyranosyl ring that were in mismatch with the modelled ligand orientation in PDB 5a1a. Refinement against the LocScale map led to re-orientation of the ligand (Figure 4G-J) and several active site residues. Although the overall ligand shifts are small (0.7 – 1.0Å, rmsd 1.468 Å over 20 atoms), the ligand conformation obtained with the LocScale map satisfies additional hydrogen bonds in the active site and results in tighter accommodation of the inhibitor in the binding pocket (Figure 4K/L).

We also generated LocScale maps for TRPV1, Pol III and γ-secretase. For both TRPV1 and Pol III, the respective LocScale maps provided enhanced contrast in regions of previously fragmented and weak density. The LocScale maps allowed the rebuilding of loop regions and side chains in the TRPV1 structure (Figure 5 A-C) and confident placement of predicted secondary structure elements in previously uninterpretable parts in the C82 subunit of Pol III (Figure 5 D-F). Interestingly, when inspecting the γ-secretase LocScale map, we found additional density contrast at 9 out of 11 N-glycosylation sites, which would permit modeling glycan extensions beyond the deposited carbohydrate chains (Figure 6A-C). Since the density did not allow unambiguous identification of chemical identity, we in these cases refrained from building additional carbohydrate residues. Notably, in addition to previously described sites, the LocScale map revealed interpretable density at four additional and previously unreported N-linked glycosylation sites that allowed building of the glycan base (Figure 6D/E). Together, these examples showcase the potential of locally enhanced contrast in uncovering weak but relevant map features, and its importance for map interpretation and model building.

**Figure 5.**
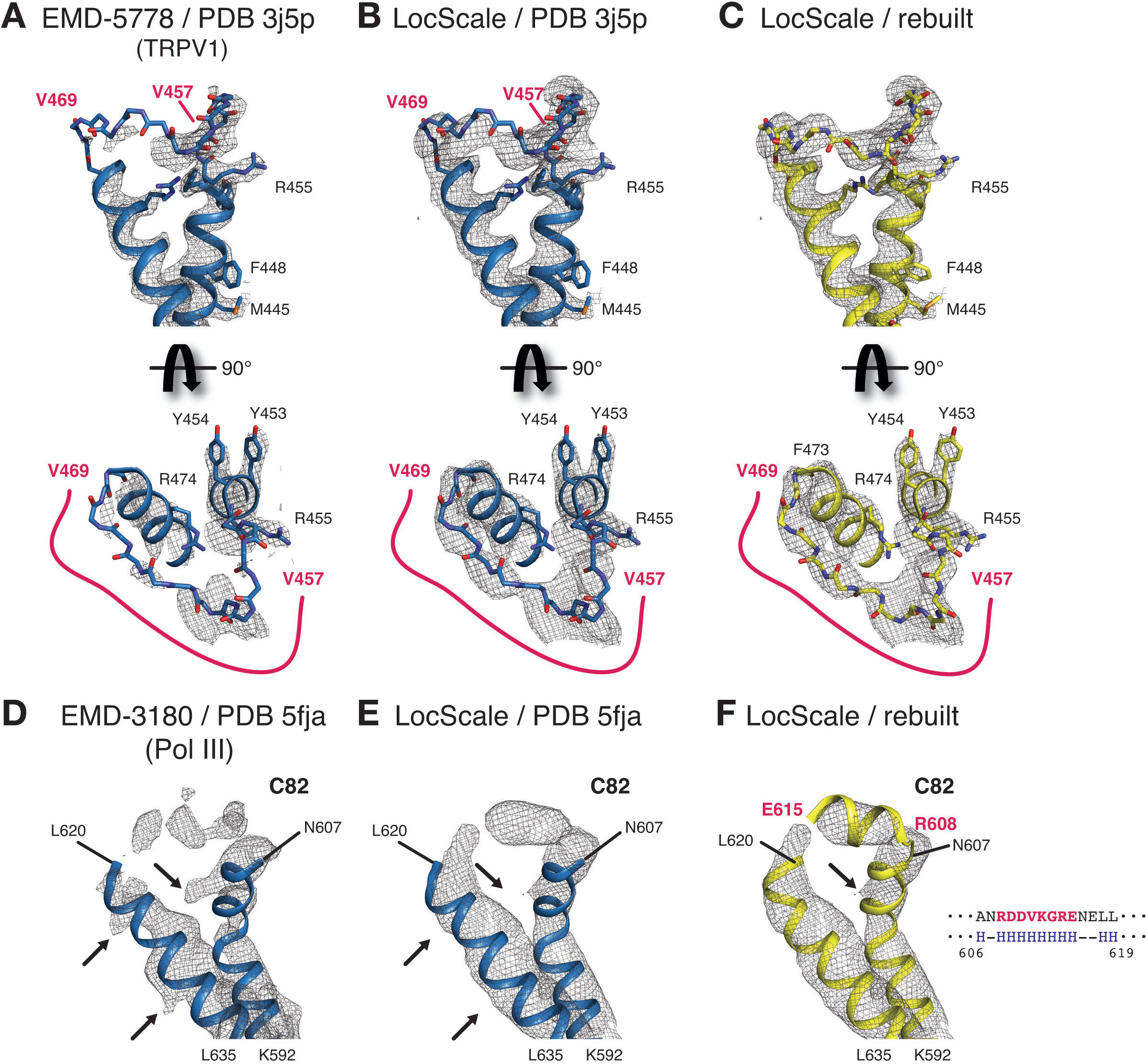
LocScale maps for TRPV1 and RNA Pol III. (**A**) Close-up density for TRPV1 channel (EMD-5778) superposed on the deposited coordinate model (PDB ID 3j5p). The map was additionally sharpened with a B-factor of −100 Å^2^ to make densities for aromatic and other large side chains more clearly visible. Transmembrane views (top) and the corresponding perpendicular view (bottom) of the V457–V469 loop region are displayed, showing poorly resolved density for V457–V469 residues (main chain shown in stick representation). Selected residues with large side chains are also shown in stick representation. (**B**) Same view as (A) with PDB ID 3j5p superposed onto the LocScale density. Density is shown at a threshold level matching visible side-chain densities in (A). Continuous density is observed for residues V457–V469, allowing less ambiguous building of the main chain trace. (**C**) LocScale density for the same regions shown in (A) and (B) superposed on the atomic model obtained after adjustment and refinement against the LocScale map. (**D**) Close-up density for Pol III subunit C82 (EMD-3180) superposed onto the deposited atomic model (PDB 5fja) shown in cartoon representation. Density for residues R608–L619 is fragmented and precludes model building. Note the pronounced density artifacts (arrows) incompatible with side chain densities. (**E**) Same view as (D) with PDB 5fja superposed onto the LocScale density. The density is suggestive of a helical conformation for residues R608-E615, confirming secondary structure predictions (right). Density artifacts from (D) are reduced in the LocScale map. (**F**) C82 model rebuilt and refined using the LocScale map. The predicted helix formed by residues R608-E615 fits the helical density as suggested by the LocScale map.

**Figure 6.**
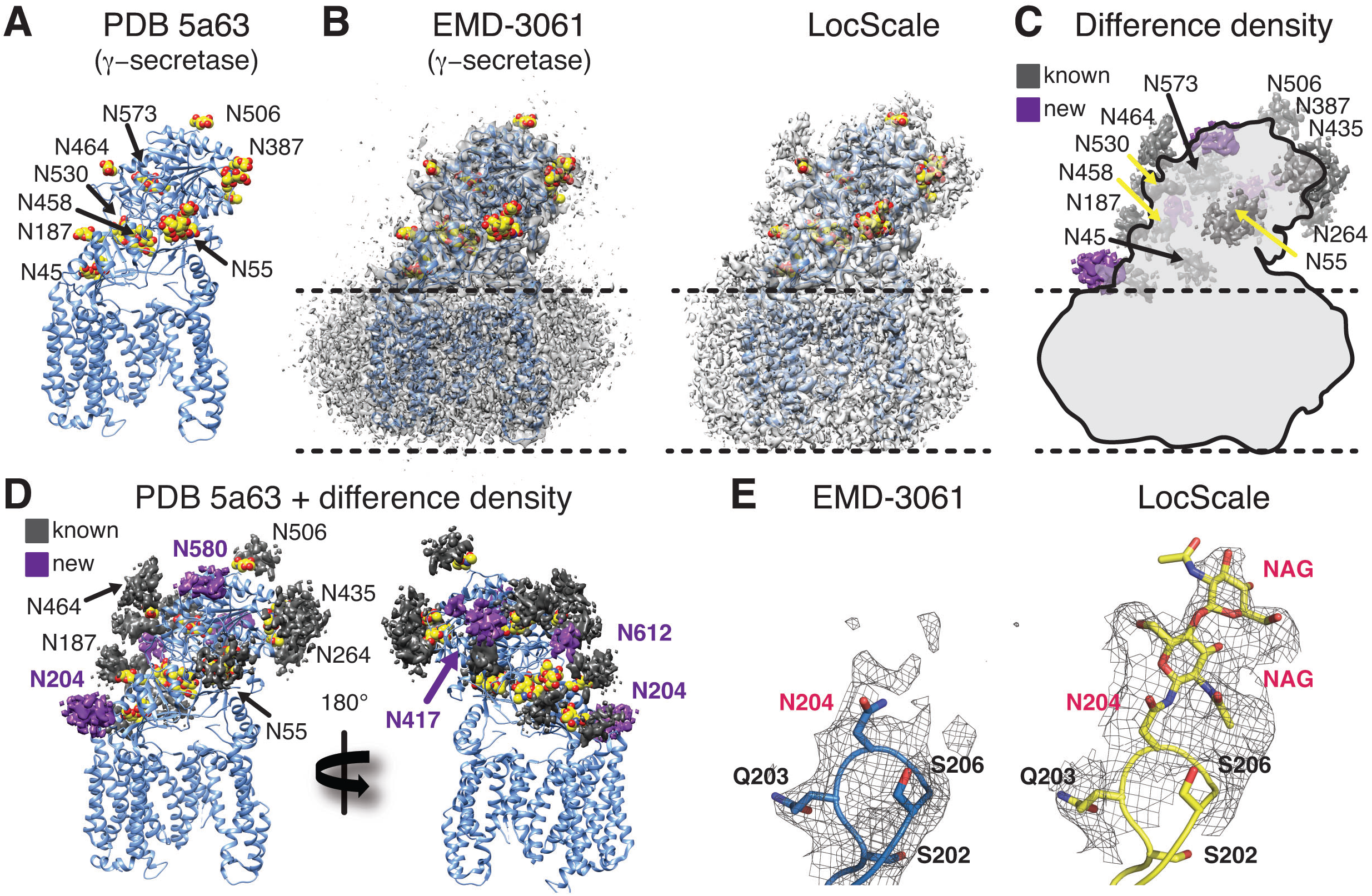
Enhanced visibility of glycan chains in LocScale maps of γ-secretase. (**A**) Atomic model of γ-secretase (PDB ID 5a63) in cartoon representation. Modeled glycan residues in nicastrin are shown as spheres and N-linked glycosylation sites are indicated. (**B**) Comparison of EMD-3061 (left) and LocScale (right) maps contoured at 3σ with ribbon model as in (A). (**C**) Difference density of (B left and right) superposed on light gray silhouette representation of EMD-3061. Regions with difference density beyond 4σ located at known glycosylation sites are colored in dark gray and newly identified glycosylation sites are shown in purple. (**D**) Difference density from (C) mapped onto the atomic model of γ-secretase. Purple difference density locates to putative additional glycosylation sites. (**E**) Density around N204 of nicastrin for EMD-3061 (left) and LocScale map (right) reveals additional density for two N-acetylglucsoamine (NAG) residues in the LocScale map.

### LocScale maps are improved map targets for model refinement

In order to assess quantitatively whether LocScale density improvement positively affects atomic model refinement, we performed coordinate refinement using either the EMDB-deposited or the LocScale map as minimization targets. Initially, we randomly perturbed the coordinates of the PDB-deposited coordinates of all four structures (β-galactosidase: PDB 5a1a; TRPV1: PDB 3j5p; Pol III: PDB 5fja; γ-secretase: PDB 5a63) and subjected the resulting models to iterative real-space refinement against the EMDB-deposited map and the corresponding LocScale map (Adams et al., 2010; Hoffmann et al., 2015). In every case, the overall real-space correlation improved by 4 – 6% for models refined against the LocScale map (Table 1), and FSC curves computed between model maps and original reconstructions show small but notable improvements of the LocScale model fit (Table 1–figure supplement 1). In particular, FSC improvements are largest for TRPV1 and Pol III, for which also the resolution differences across the map are largest. In addition, we find improved EMRinger and Molprobity scores (Barad et al., 2015; Chen et al., 2010) for all models refined with LocScale maps, indicating that better density fit is not achieved at the expense of model distortions.

**Table 1.** Model refinement statistics.

|  | **β-galactosidase<sup>1</sup>** |  | **TRPV1 Channel<sup>2</sup>** |  | **RNA polymerase III<sup>3</sup>** |  | **γ-secretase<sup>4</sup>** |  |
| --- | --- | --- | --- | --- | --- | --- | --- | --- |
|  | EMD-2984 | LocScale | EMD-5778 | LocScale | EMD-3180 | LocScale | EMD-3061 | LocScale |
| Starting model PDB ID | 5a1a |  | 3j5p |  | 5fja |  | 5a63 |  |
| Resolution (Å)<br>LocRes range (Å) | **2.2**<br>(1.9 - 3.6) |  | **3.4**<br>(2.5 - 4.8) |  | **4.7**<br>(3.4 - 8.1) |  | **3.4**<br>(3.0 - 6.8) |  |
| Refinement (Å) | 186.0 – 1.9 |  | 311.3 – 2.5 |  | 320.9 – 3.4 |  | 252.0 – 3.0 |  |
| Overall RSCC* | 0.70 | 0.75 | 0.62 (0.75)<sup>##</sup> | 0.67 (0.81)<sup>##</sup> | 0.70 | 0.76 | 0.71 | 0.77 |
| EMRinger score | 4.03 | 4.90 | 0.24 | 0.77 | 0.50** | 1.31** | 2.60 | 2.87 |
| MOLPROBITY score | 2.04 | 1.86 | 2.69 (1.44)<sup>##</sup> | 1.98 (1.35) | 2.68 | 2.24 | 2.01 | 1.81 |
| Clash score<sup>#</sup> (all atoms) | 6.63 | 3.14 | 6.88 (2.42)<sup>##</sup> | 5.29 (2.09)<sup>##</sup> | 16.44 | 11.04 | 6.44 | 5.89 |
| Rotamer outliers (%) | 3.32 | 4.92 | 16.38 (0.00)<sup>##</sup> | 2.23 (0.00)<sup>##</sup> | 2.53 | 1.48 | 1.53 | 0.86 |
| Ramachandran statistics |  |  |  |  |  |  |  |  |
| Favored (%) | 95.98 | 96.57 | 94.3 (93.6)<sup>##</sup> | 93.4 (94.6)<sup>##</sup> | 83.2 | 85.4 | 92.86 | 91.74 |
| Disallowed (%) | 0.00 | 0.00 | 0.35 (0.64)<sup>##</sup> | 0.17 (0.64)<sup>##</sup> | 1.09 | 0.99 | 0.33 | 0.33 |
| RMS bonds (Å) | 0.008 | 0.01 | 0.008 | 0.006 | 0.008 | 0.01 | 0.011 | 0.012 |
| RMS angles (°) | 1.41 | 1.50 | 1.71 | 1.44 | 1.67 | 1.32 | 1.41 | 1.53 |

\* Overall real-space correlation computed at the average map resolution using a soft mask around atoms. The EMDB deposition was used as the reference map in all cases.
\*\* EMRinger score calculated only for parts of structure with local resolution estimate better than 4.5 Å.

To better discriminate the individual effects of low-pass filtration and local contrast enhancement on real-space refinement targets, we also examined uniformly sharpened, local resolution-filtered (LocRes) maps (see Methods) as targets for atomic model refinement. We observed that LocRes maps partially compensate for artifacts arising from uniform sharpening in 3D reconstructions with significant resolution variation, whereas local contrast optimization in LocScale maps consistently provides additional detail (Table1–figure supplement 2). Comparison of global refinement statistics using LocRes and LocScale maps as targets support these qualitative observations (Table 1–table supplement 1). Together, the results from four examples show that iterative local contrast optimization by LocScale density improvement results in atomic models with increased overall density fit and improved stereochemistry.

## Discussion

We have introduced a procedure that iteratively improves the interpretability of cryo-EM density maps. The LocScale method uses prior information from an atomic reference model that itself improves progressively during rounds of alternating coordinate refinement and density improvement. Our method is generally applicable to any cryo-EM structure whose density can be interpreted in terms of an atomic model and thus only requires the input of density map and superimposed atomic coordinates without further specification of tuning parameters. Current contrast enhancing approaches often inadvertently result in either omission of structural detail or amplification of background noise, resulting in map artifacts that often cannot be easily distinguished from real atomic features. By applying amplitude scaling locally, LocScale maps represent high-resolution signal more accurately throughout the map (Figures 3-6). The procedure is reminiscent of an optimized low-pass filter that adaptively scales and filters the map amplitudes to match the local resolution and expected mass density distribution of the underlying molecular structure. Instead of reference-based scaling as implemented in the LocScale procedure, local sharpening could also be performed after estimating the amplitude fall-off within each density window using Guinier analysis. While this approach remains an option in the absence of an atomic model, it requires the presence of sufficiently high-resolution data for reliable exponential fitting of B-factors (DeLaBarre and Brunger, 2006). Strictly, exponential sharpening is an approximation that assumes scattering to arise from randomly distributed free atoms (Wilson, 1942). One advantage of LocScale amplitude scaling over other sharpening methods is that it explicitly accounts for the characteristic local profile of the amplitude fall-off that is due to specific structural properties of secondary structure motifs (Morris et al., 2004) and can differ markedly from a simple exponential decay (Figure 2F, Figure 2 – supplementary figure 1).

Using four examples, we showed that iterative LocScale contrast restoration consistently improves interpretability of density maps and accuracy of atomic models over a range of resolutions between 2 and 8 Å. While we have only tested the utility of such maps for real-space refinement, we expect LocScale maps to equally benefit phase-restrained refinement in reciprocal space as implemented in CNS (Brunger et al., 1998) or REFMAC (Murshudov et al., 1997). Although we refer to LocScale as a form of density improvement, it is important to distinguish our procedure from what is commonly understood by density modification in X-ray crystallographic refinement. Crystallographic density modification improves map quality through obtaining new phase estimates by modification of the electron density to match expected properties such as flatness or disorder of the solvent region (Cowtan, 2010; B. C. Wang, 1985; Zhang et al., 1997). In the LocScale approach put forward here, density modification refers to improvements of the amplitude component of the molecular structure factor such that it conforms to prior expectations from a reference model, whereas the phase component is retrieved directly from the experiment and remains unchanged by the procedure. As the method is restricted to scaling of amplitudes against a radial average, we argue that reference bias from the atomic model is not present. We show that for cases with incomplete models or minor conformational errors in the model, the information stored in the radial amplitude profile still provides a good initial estimate of the mass density distribution and is sufficient to restore low contrast map features more clearly when compared with overall sharpening procedures (Figure 2–figure supplement 1A-D). We find that either uniformly sharpened or globally scaled maps, optionally combined with filtering based on local resolution estimates, are appropriate starting targets for initial model building and refinement of atomic coordinates and B-factors. Once estimates of atomic B-factors have been obtained, LocScale maps provide locally optimized contrast and facilitate further rounds of model building and refinement. While it has been shown that map sharpening positively affects model refinement (Afonine et al., 2015; Joseph et al., 2016), a recent report suggested that overall benefits are limited (R. Y.-R. Wang et al., 2016). In our test cases with different ranges of resolution, we do confirm previously described benefits of sharpening and observe a notable improvement of refined models using LocScale maps, likely because locally optimized sharpening levels result in improvements of density fit and geometry over all map areas.

Regardless of resolution, cryo-EM maps degrade in quality towards the particle periphery due to limitations of current alignment and 3D reconstruction algorithms. Furthermore, true structural flexibility is expected to be more prominent in the particle periphery as plunge freezing captures the solution-state conformational landscape at the respective experimental condition. While part of this conformational heterogeneity can be dealt with in unsupervised 2D and 3D image classification procedures (Jonic, 2017; Lyumkis et al., 2013; Scheres, 2012), remaining flexibility degrades resolution and results in localized blurring of the density map. The need for approaches that address resolution variation has been recognized and local resolution filtering has been proposed (Cardone et al., 2013; Fernandez-Leiro and Scheres, 2016; Moriya et al., 2017). The LocScale procedure takes the concept of local filtering further by optimizing contrast of the cryo-EM density through local amplitude scaling against an atomic reference model. In the four presented test cases, density improvements were particularly apparent in regions of increased structural flexibility such as peripheral subunits or flexible loop regions, as well as partial occupancy sites of ligands, active site residues and carbohydrate chains at N-linked glycosylation sites. It is not uncommon that such parts of protein complexes contain important information about the inhibition mechanism and interaction interfaces. Atomic model refinement followed by a local amplitude scaling step results in a single map that displays structural information optimally for all map areas simultaneously, including those parts with large resolution variation.

For best performance, LocScale density improvement requires good estimates of atomic B-factors associated with the reference coordinates. We show, however, that density modification is sufficiently robust to result in improved maps even with reference structures for which atomic B-factors have not yet been refined. Refinement of B-factors at low resolution is challenging and includes the risk of over-fitting as the number of experimental observables is typically much lower than the number of parameters to be fitted (DeLaBarre and Brunger, 2006). Considering that atomic B-factors represent the square amplitude of displacement from the refined coordinates they correlate with resolution. We thus propose to use the agreement of B-factor and local resolution estimated on a per-residue basis as a low-level assessment tool for the robustness of the atomic B-factor refinement (**Figure 2–figure supplement 2B**). Finding better ways to refine and validate B-factors at low resolution in real and reciprocal space remains an important area of investigation with potential for further improvements in X-ray crystallographic as well as in cryo-EM model refinement (Brunger et al., 2009). In the test cases investigated here, we found atomic B-factor refinement to work acceptably up to a resolution of ∼8Å. Therefore, we expect the iterative LocScale density improvement to work best for near-atomic resolution cryo-EM reconstructions that are appropriate for model building and allow reliable coordinate refinement including B-factors. Interestingly, inspection of FSC_ref_ curves often displays notable disagreement between FSC_ref_ and half-map FSCs at low resolution despite very good real-space density correlations between the model and density map, indicating that additional efforts are required to accurately model molecular structure factors including solvent contributions (Clarage et al., 1992; Jiang and Brunger, 1994); and the need for inclusion of more sophisticated B-factor models to reflect concerted molecular motion of atom groups in cryo-EM model refinement strategies (Painter and Merritt, 2006).

For the future of cryo-EM, it is tempting to speculate that atomic models will become more tightly integrated into the structure determination process. In principle, up-stream parts of the structure determination process such as the orientation search of the particle images for 3D reconstruction may also benefit from efforts that aim at iteratively improving both density map and atomic model, analogous to X-ray crystallography procedures that refine the atomic model against diffraction data. We here put forward an iterative cryo-EM density improvement that makes use of local amplitude scaling derived from a refined atomic reference model while avoiding bias from that model. Cryo-EM density improvement by the LocScale procedure results in more faithful density representations and at the same time overcomes previous shortcomings in dealing with maps displaying considerable resolution variation. The approach is expected to be beneficial for a wide range of cryo-EM structures and will enhance atomic model interpretation of the rapidly growing number of high-resolution cryo-EM density maps.

## Methods

### Implementation

All computational methods have been written in Python (www.python.org) and make use of libraries and functions from SPARX/EMAN2 (Hohn et al., 2007) and CCTBX (Adams et al., 2010). Numerical operations are performed using the Numpy package and Scipy libraries (www.scipy.org). The program LOCSCALE and associated scripts including instructions will become available for download upon acceptance of the manuscript from the authors’ website.

### Model-based amplitude scaling (LocScale maps)

The test image used in Figure 2A-E is in the public domain and the amplitude manipulations were implemented in Numpy and Scipy (www.numpy.org). For local amplitude scaling of cryo-EM densities, model maps were obtained from atomic coordinates by computing B-factor weighted structure factors using electron atomic form (Colliex et al., 2006) and their conversion into density by inverse Fourier transform using a CCTBX-based routine (pdb2map.py). LOCSCALE accepts unfiltered and unsharpened volumes, i.e. untreated 3D reconstructions, for scaling. Experimental and model maps were sampled in a rolling window of (n × n × n) voxels. For each grid step, the unfiltered experimental density map contained within the rolling window was sharpened by a resolution-dependent amplitude scaling factor using

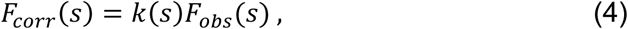
where *F*_*obs*_(*s*) is the Fourier amplitude of the original density map at spatial frequency *s*, and

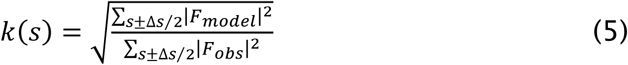
is the scale factor at frequency *s* derived from the ratio of radially averaged power spectra integrated within a Fourier shell of thickness Δ*s*. *F*_*model*_(*s*) and *F_obs_*(*s*) indicate Fourier amplitudes from model and experimental map, respectively. The map value of the central voxel contained in the scaled volume element is then assigned to the corresponding voxel of the new map. Rolling windows were padded by two times the window size to increase precision in Fourier space. Repeating the procedure over each voxel position yields a map representation with optimized contrast.

The program accepts an unfiltered 3D reconstruction and a superposed atomic model as input. Alternatively, a precomputed model map can also be provided alongside the unfiltered 3D reconstruction. A soft-edged mask can be supplied to restrict the computations to the voxels contained within the mask. The program requires *dmin* and *window_size* as additional input parameters. The parameter *dmin* refers to the resolution to which the reference map should be simulated and scaling should be performed (typically Nyquist frequency) and *window_size* denotes the pixel length of the density cube contained in the rolling window. As a typical window size parameter, we used seven times the resolution in pixels in an effort to balance small window size to preserve locality of the approach against large window size to have sufficient number of Fourier pixels to appropriately sample the fine-structure of the amplitude fall-off.

### Atomic model refinement

Unfiltered reconstructions for TRPV1 (EMD-5778), γ-secretase (EMD-3061) and β-galactosidase (EMD-2984) were obtained from the EMDB model challenge website (http://challenges.emdatabank.org). Initial map targets for coordinate refinement were generated by applying uniform filtering to the original EMDB entries. The deposited map for TRPV1 (EMD-5778) was globally sharpened using an additional B-factor of −100 Å^2^. Coordinate refinement was performed using real-space refinement against the initial map as implemented in cctbx/PHENIX (Adams et al., 2010) with additional restraints on secondary structure. Residue-grouped atomic B-factors were refined in reciprocal space. For TRPV1 and β-galactosidase, non-crystallography symmetry (NCS) restraints were employed to account for rotational and dihedral symmetry. Reference restraints for PETG were obtained with *phenix.elbow* from the crystal structure of 2-phenylethyl 1-thio-β-D-galactopyranoside (Brito et al., 2011). Building of glycans used glycan modeling tools available in Coot. Local resolution for all maps was estimated by local FSC computations using an in-house Python program *locres.py*. Local resolution was mapped to the coordinate models using the *measure_mapValues* function in UCSF Chimera (Pettersen et al., 2004), and the residue-averaged resolution was used to assign local resolution-scaled reference restraints. Model rebuilding was performed in Coot (Emsley and Cowtan, 2004). Half-set reconstructions for cross validation were obtained from the EMDB model challenge. Model maps were generated by inverse Fourier transform using B-factor-weighted electron form factors (Colliex et al., 2006). The FSC_ref_ between model map and half-set 3D reconstructions was used to assess over-fitting. To this end, coordinates were first refined against the full map. Coordinates were then randomly displaced by a maximum of 0.5 Å and subjected to three cycles of real-space refinement against one of the half maps (work map) using the same protocol outlined above. The other half map (test map) was used as the test map for cross-validation. Following refinement, we computed the FSC of the refined model against the work map (FSC_work_) and the cross-validated FSC between refined model and the test map (FSC_test_). All FSCs were computed using a structure mask obtained from the respective EMDB entry that was low-passed filtered to 60 Å.

To compare model refinement against deposited EMDB entries and LocScale maps, PDB-deposited coordinate models associated with the respective EMDB entry were randomly perturbed by applying atom shifts of up to 0.4 Å to serve as starting models. Five iterations of local and global real-space coordinate refinement against the respective EMDB or LocScale map were each followed by refinement of atomic B-factors in reciprocal space. As above, secondary structure restraints, resolution-dependent weights and NCS restraints were employed. EMRinger scores were computed using *phenix.em_ringer* (Barad et al., 2015); all other validation scores were obtained using MOLPROBITY (Chen et al., 2010). All cross-validation FSC and real-space correlations were computed against the original reconstruction and the deposited EMDB map, respectively.

## Acknowledgements

AJJ acknowledges financial support by an EMBL Interdisciplinary Postdoc (EIPOD) fellowship under Marie Curie Actions (PCOFUND-GA-2008-229597), a Marie-Sklodowska-Curie IEF fellowship (PIEF-GA-2012-331285), the Deutsche Forschungsgemeinschaft (DFG) through the excellence cluster “The Hamburg Center for Ultrafast Imaging (CUI) – Structure, Dynamics and Control of Matter at the Atomic Scale” (EXC1074) and the Joachim Herz Foundation.

---

**Figure 2–figure supplement 1.**
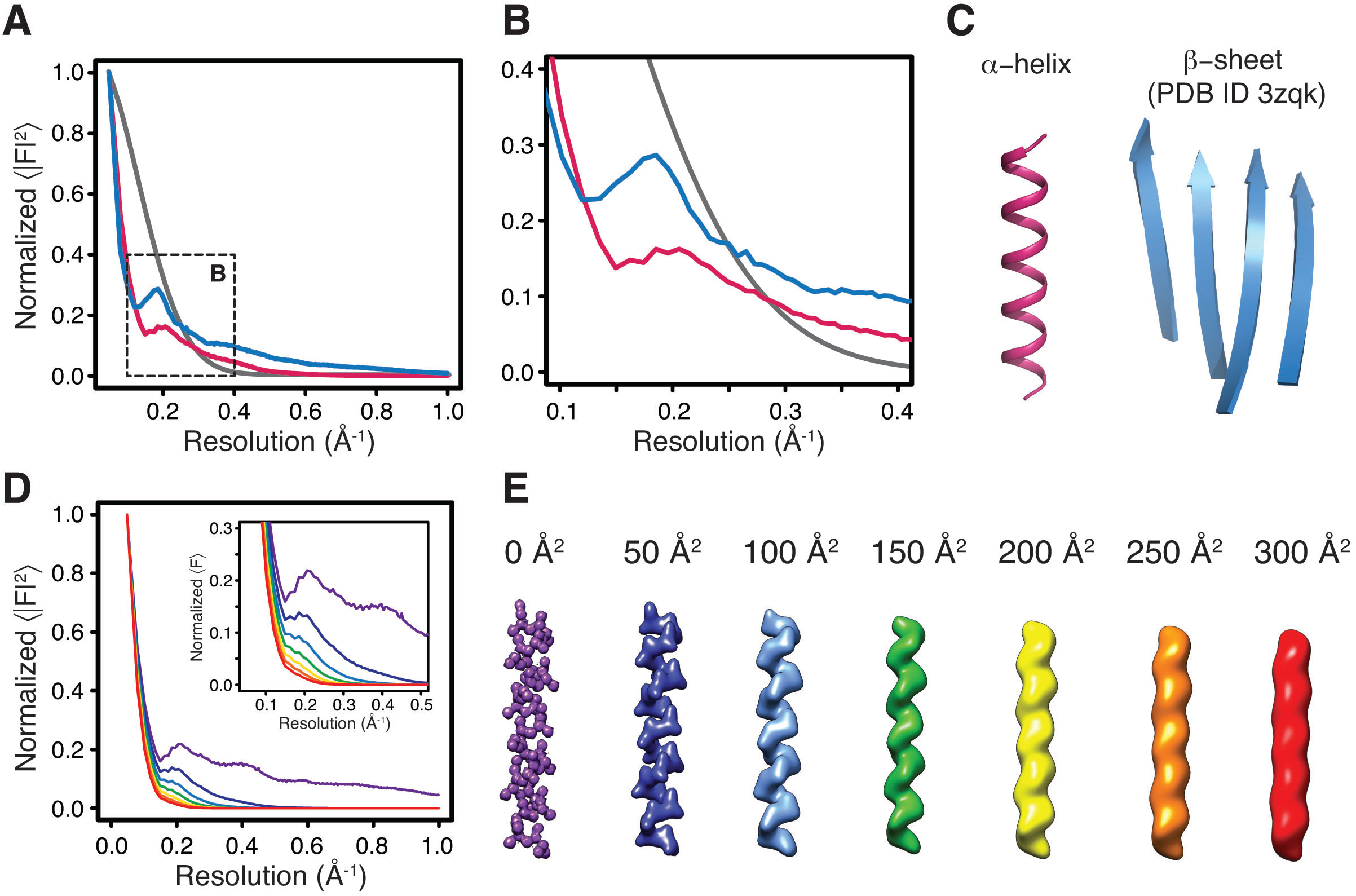
Effect of secondary structure and atomic B-factors on the radial structure factor amplitude profile. (**A**) Typical amplitude profiles (magnified in (**B**)) computed from density maps simulated at 1 Å resolution of an idealized α-helix (pink), a representative β-sheet from PDB ID 3zqk (blue) and a random distribution of atoms (grey) (**C**). Note the characteristic differences of the profiles of α-helical and β-sheet structures in comparison with the exponential decay from randomly distributed atoms. (**D**) Illustration on the amplitude profile dependence on the atomic B-factor for the density of an idealized α-helix (**E**). Note the gradual disappearance of the characteristic profile at high B-factors. Color-coding for profiles at different B-factors follows that in (E).

**Figure 3–figure supplement 1.**
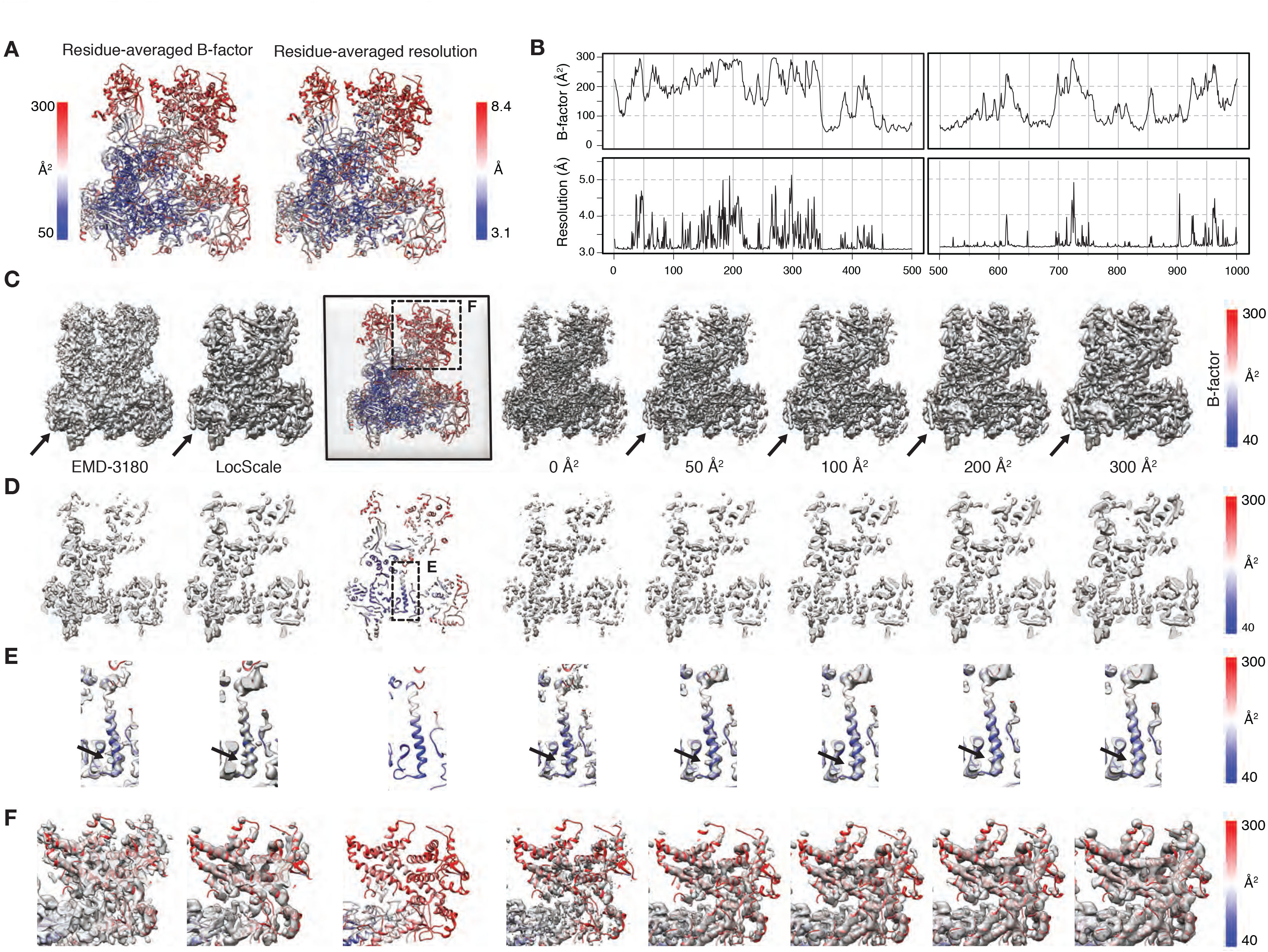
Effect of atomic B-factors on LocScale procedure. **(A)** Comparison of residue-averaged B-factor (left) and local map resolution (right) mapped onto a cartoon representation of Pol III. The spatial distribution of atomic B-factors follows that of local resolution, with low B-factors (high resolution) in the core and high B-factors (low resolution) at the periphery. **(B)** Stacked diagrams of atomic B-factors (top) and local resolution (bottom) for two residue stretches of Pol III. Both quantities show correlating profiles of maxima and minima. **(C)** Effect of atomic B-factors on the LocScale procedure. The atomic model (PDB ID 5fja) is shown in cartoon representation with residue-averaged B-factors following the coloring scheme from (A). The deposited map (EMD-3180) and final LocScale map calculated using a reference map simulated from PDB ID 5fja are shown on the left. Right to the atomic model we show LocScale maps obtained with reference maps simulated from a coordinate model with uniform isotropic atomic B-factors of 0, 50, 100, 200 and 300 Å^2^, respectively. Each map element resolved at a particular resolution has a corresponding B-factor that best restores contrast in the respective LocScale map. **(D)** Central density slice for the presented densities in (C). **(E)** Close-up of a central helix with superimposed density. Note the progressive disappearance of side chain features in the LocScale maps (arrow) with atomic B-factors that are too high (≥200 Å^2^). **(F)** Close-up of a poorly resolved heterodimer region of Pol III as indicated in (C). When the average B-factor is too low (B=0 Å^2^), peripheral helix map features become weak or fragmented. High B-factors (B=300 Å^2^) lead to the appropriate low-pass filtered appearance of the same peripheral helix.

**Figure 4–figure supplement 1.**
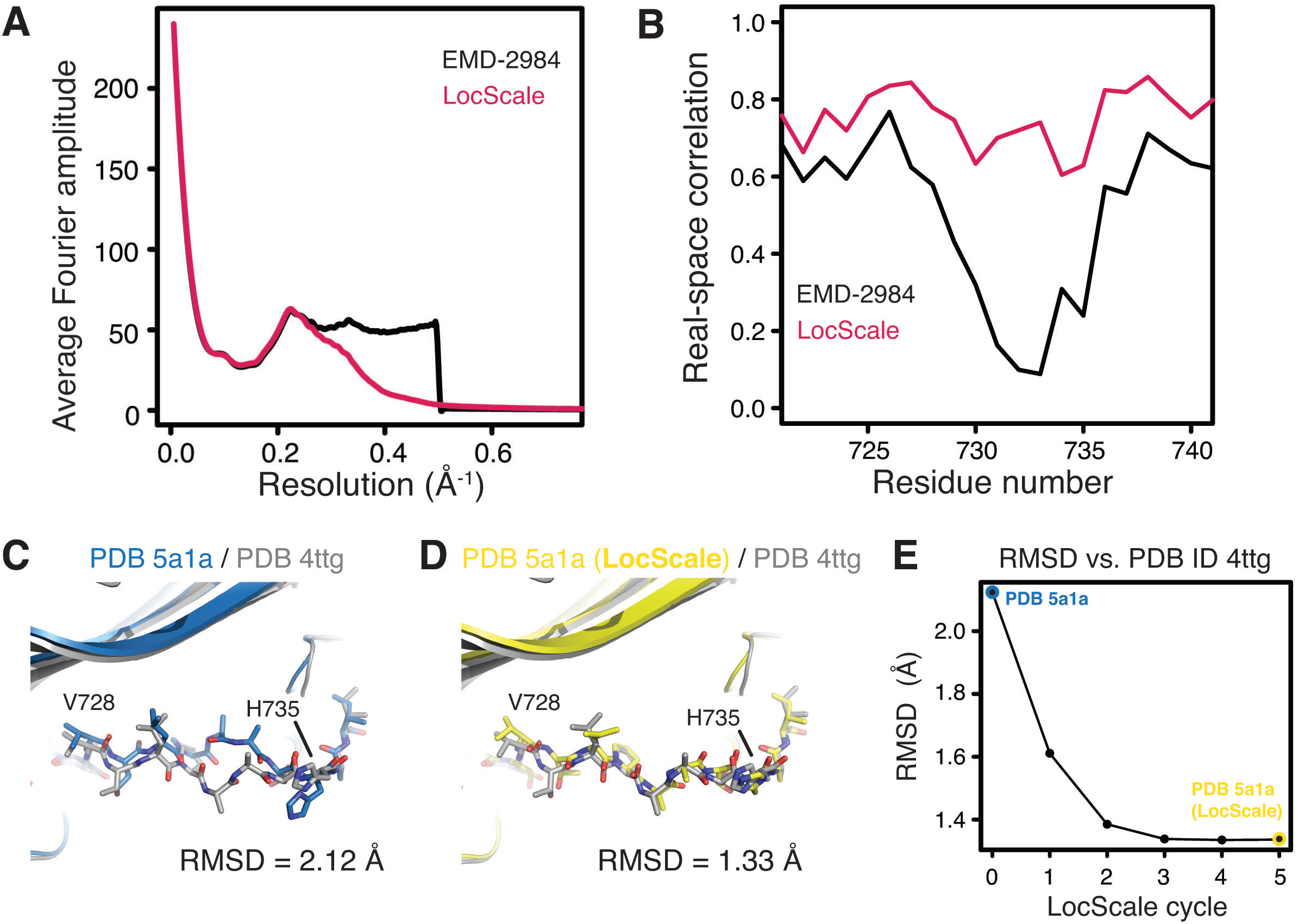
Comparison of β-galactosidase radial amplitude profiles and iterative refinement of coordinates from the V728–H735 loop region. **(A)** Radial amplitude spectrum of LocScale β-galactosidase map. In comparison with the deposited map obtained by uniform B-factor sharpening, the LocScale procedure results in a smooth fall-off to high-resolution amplitudes. **(B)** Residue-based real-space correlation improves in the V728–H735 loop region for the conformation built using the LocScale map. (**C**) Comparison of the V728–H735 stretch of PDB ID 5a1a (blue) with the 1.6 Å crystal structure of *E.coli* β-galactosidase PDB ID 4ttg (silver). The RMSD states the root mean square deviation between PDB IDs 5a1a and 4ttg computed across all atoms in the sequence V728–H735. (**D**) Same view as (C) comparing the model after rebuilding and iterative refinement using the LocScale procedure (yellow). (**E**) Plot of the model differences in RMSD (Å) illustrated in (C) and (D) as a function of alternating LocScale and refinement cycle.

**Table 1–figure supplement 1.**
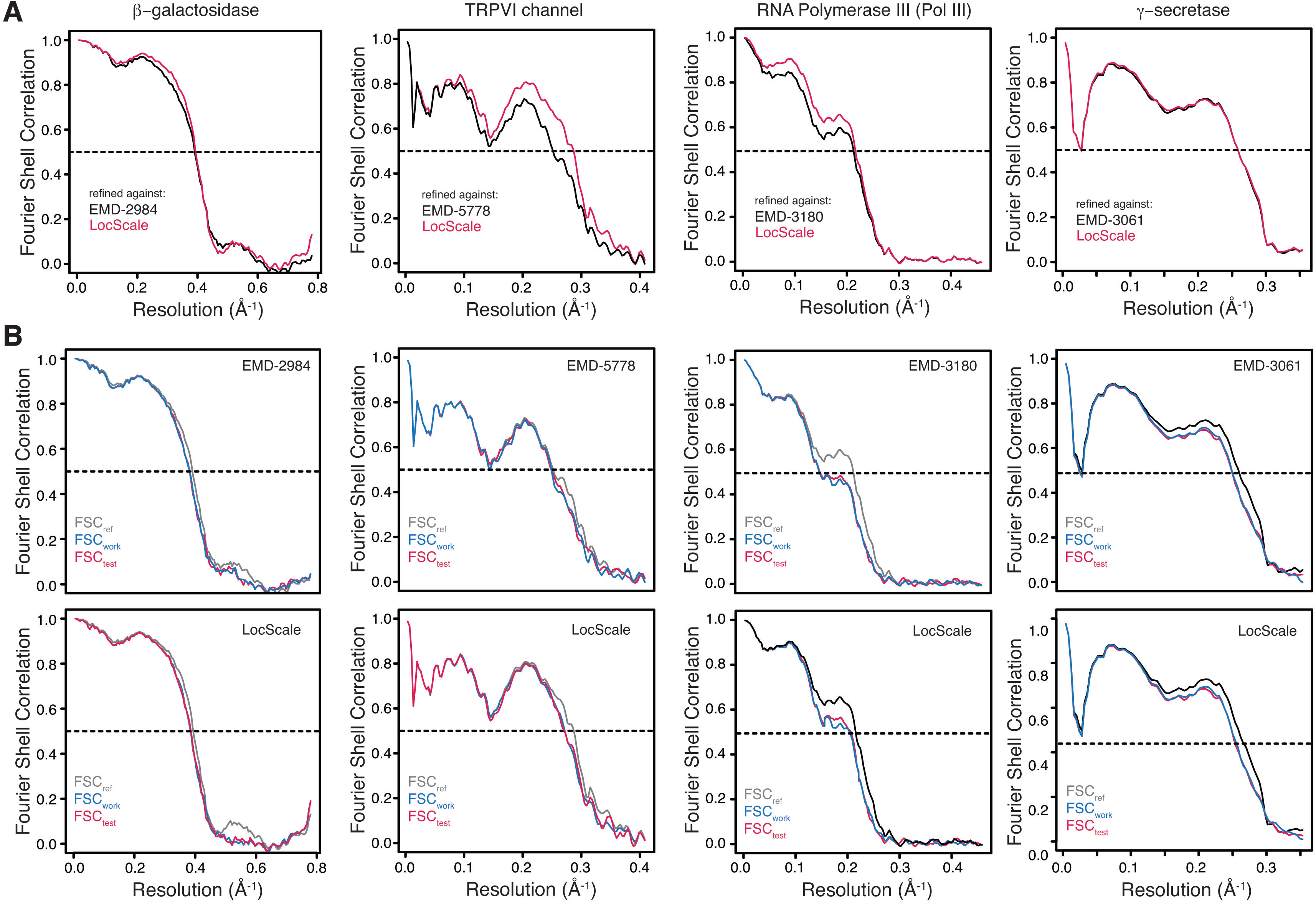
Model refinement statistics for local resolution-filtered maps. (**A**) FSC_ref_ curves for β-galactosidase, TRPV1, Pol III and γ-secretase models refined against the original EMDB entry (black) or the respective LocScale map (red). All FSC_ref_ curves were computed using the unfiltered and unsharpened original reconstruction. (**B**) FSC curves for work (FSC_work_, blue) and cross-validated test maps (FSC_test_, red) from model refinement and validation using independent half maps shown for both refinement against the original PDB entry (top) and the respective LocScale maps (bottom). The closely matching profiles of FSC_work_ and FSC_test_ indicates that no significant overfitting took place. The corresponding FSC_ref_ (gray) are shown for comparison.

**Table 1–figure supplement 2.**
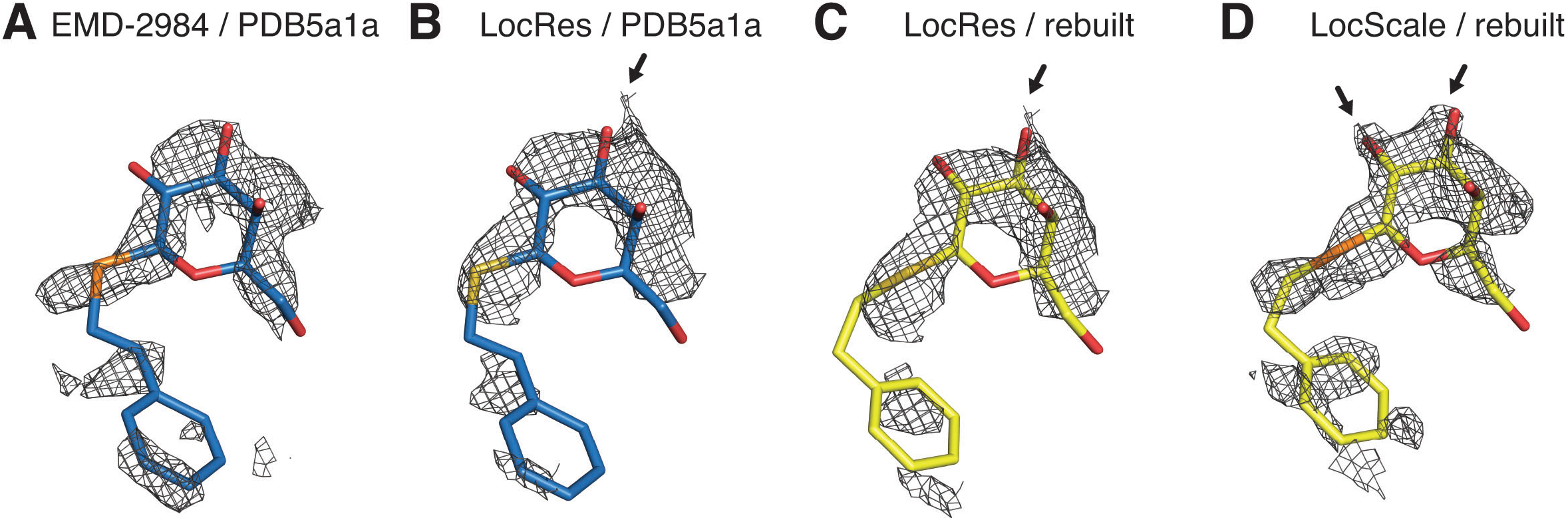
LocScale maps recover contrast optimally when compared with local resolution-filtered (LocRes) maps. (**A**) Density for the PETG ligand bound in EMD-2984 superposed on the PETG conformation from PDB ID 5fja shown in stick representation (replicate from Figure 4G). All densities are carved around the PETG ligand with a 1.6 Å carve radius. (**B-C)** PETG from PDB ID 5fja (B) or the model obtained after refinement against the LocScale map (C) superposed on the density of EMD-2984 filtered according to local resolution (LocRes). (**D**) LocScale density with PETG ligand as in Figure 4J. Only the LocScale map recovers density contrast optimally to reveal the hydroxyl substituents of the hexa-pyranosyl ring.

**Table 1–figure supplement 1.** Model refinement statistics for local resolution-filtered maps.

| | β-galactosidase | TRPV1 Channel | RNA polymerase III | γ-secretase |
| --- | --- | --- | --- | --- |
|  | LocRes | LocRes | LocRes | LocRes |
| Starting model PDB ID | 5a1a | 3j5p | 5fja | 5a63 |
| Resolution (Å)<br>LocRes range (Å) | **2.2**<br>(1.9 - 3.6) | **3.4**<br>(2.5 - 4.8) | **4.7**<br>(3.4 - 8.1) | **3.4**<br>(3.0 - 6.8) |
| Refinement (Å) | 186.0 – 1.9 | 311.3 – 2.5 | 320.9 – 3.4 | 252.0 – 3.0 |
| Overall RSCC* | 0.72 | 0.64 (0.78)<sup>##</sup> | 0.72 | 0.73 |
| EMRinger score | 4.15 | 0.46 | 0.89** | 2.69 |
| MOLPROBITY score | 2.00 | 2.43 (1.40)<sup>##</sup> | 2.62 | 1.95 |
| Clash score<sup>#</sup> (all atoms) | 4.45 | 6.45 (2.34)<sup>##</sup> | 13.31 | 6.18 |
| Rotamer outliers (%) | 3.83 | 4.63 (0.00)<sup>##</sup> | 1.95 | 1.00 |
| Ramachandran statistics |  |  |  |  |
| *Favored* (%) | 96.24 | 95.1 (94.9) | 86.4 | 93.01 |
| *Disallowed* (%) | 0.00 | 0.31 (0.64)<sup>##</sup> | 1.00 | 0.48 |
| RMS bonds (Å) | 0.008 | 0.008 | 0.01 | 0.01 |
| RMS angles (°) | 1.48 | 1.43 | 1.42 | 1.50 |

## References

Adams, P.D., Afonine, P.V., Bunkóczi, G., Chen, V.B., Davis, I.W., Echols, N., Headd, J.J., Hung, L.-W., Kapral, G.J., Grosse-Kunstleve, R.W., McCoy, A.J., Moriarty, N.W., Oeffner, R., Read, R.J., Richardson, D.C., Richardson, J.S., Terwilliger, T.C., Zwart, P.H., 2010. PHENIX: a comprehensive Python-based system for macromolecular structure solution. Acta Crystallogr D Biol Crystallogr 66, 213–221. doi:10.1107/S0907444909052925

Adrian, M., Dubochet, J., Lepault, J., McDowall, A.W., 1984. Cryo-electron microscopy of viruses. Nature 308, 32–36.

Afonine, P.V., Moriarty, N.W., Mustyakimov, M., Sobolev, O.V., Terwilliger, T.C., Turk, D., Urzhumtsev, A., Adams, P.D., 2015. FEM: feature-enhanced map. Acta Crystallogr D Biol Crystallogr 71, 646–666. doi:10.1107/S1399004714028132

Agarwal, R.C., Isaacs, N.W., 1977. Method for obtaining a high resolution protein map starting from a low resolution map. Proceedings of the National Academy of Sciences 74, 2835–2839.

Allegretti, M., Mills, D.J., McMullan, G., Kühlbrandt, W., Vonck, J., 2014. Atomic model of the F420-reducing [NiFe] hydrogenase by electron cryo-microscopy using a direct electron detector. eLife 3, e01963–e01963. doi:10.7554/eLife.01963.025

Amunts, A., Brown, A., Bai, X.C., Llacer, J.L., Hussain, T., Emsley, P., Long, F., Murshudov, G., Scheres, S.H.W., Ramakrishnan, V., 2014. Structure of the Yeast Mitochondrial Large Ribosomal Subunit. Science 343, 1485–1489. doi:10.1126/science.1249410

Bai, X.-C., Fernandez, I.S., McMullan, G., Scheres, S.H.W., 2013. Ribosome structures to near-atomic resolution from thirty thousand cryo-EM particles. eLife 2, e00461. doi:10.7554/eLife.00461

Bai, X.-C., Yan, C., Yang, G., Lu, P., Ma, D., Sun, L., Zhou, R., Scheres, S.H.W., Shi, Y., 2015. An atomic structure of human γ-secretase. Nature 525, 212–217. doi:10.1038/nature14892

Barad, B.A., Echols, N., Wang, R.Y.-R., Cheng, Y., DiMaio, F., Adams, P.D., Fraser, J.S., 2015. EMRinger: side chain–directed model and map validation for 3D cryo-electron microscopy. Nat Methods 12, 943–946. doi:10.1038/nmeth.3541

Bartesaghi, A., Merk, A., Banerjee, S., Matthies, D., Wu, X., Milne, J.L.S., Subramaniam, S., 2015. Electron microscopy. 2.2 Å resolution cryo-EM structure of β-galactosidase in complex with a cell-permeant inhibitor. Science 348, 1147–1151. doi:10.1126/science.aab1576

Brito, I., Szilágyi, L., Kumar, A.A., Albanez, J., Bolte, M., 2011. 2-Phenylethyl 1-thio-β-D-galactopyranoside hemihydrate. Acta Crystallographica Section E Structure Reports Online 67, o2308–o2309. doi:10.1107/S1600536811031667

Brown, A., Long, F., Nicholls, R.A., Toots, J., Emsley, P., Murshudov, G., IUCr, 2015. Tools for macromolecular model building and refinement into electron cryo-microscopy reconstructions. Acta Crystallogr D Biol Crystallogr 71, 136–153. doi:10.1107/S1399004714021683

Brunger, A.T., Adams, P.D., Clore, G.M., DeLano, W.L., Gros, P., Grosse-Kunstleve, R.W., Jiang, J.S., Kuszewski, J., Nilges, M., Pannu, N.S., Read, R.J., Rice, L.M., Simonson, T., Warren, G.L., 1998. Crystallography & NMR System: A New Software Suite for Macromolecular Structure Determination. Acta Crystallogr D Biol Crystallogr 54, 905–921. doi:10.1107/s0907444998003254

Brunger, A.T., DeLaBarre, B., Davies, J.M., Weis, W.I., 2009. X-ray structure determination at low resolution. Acta Crystallogr D Biol Crystallogr 65, 128–133. doi:10.1107/S0907444908043795

Cardone, G., Heymann, J.B., Steven, A.C., 2013. One number does not fit all: mapping local variations in resolution in cryo-EM reconstructions. J Struct Biol 184, 226–236. doi:10.1016/j.jsb.2013.08.002

Chen, V.B., Arendall, W.B., Headd, J.J., Keedy, D.A., Immormino, R.M., Kapral, G.J., Murray, L.W., Richardson, J.S., Richardson, D.C., 2010. MolProbity: all-atom structure validation for macromolecular crystallography. Acta Crystallogr D Biol Crystallogr 66, 12–21. doi:10.1107/S0907444909042073

Cheng, Y., 2015. Single-Particle Cryo-EM at Crystallographic Resolution. Cell 161, 450–457. doi:10.1016/j.cell.2015.03.049

Clarage, J.B., Clarage, M.S., Phillips, W.C., Sweet, R.M., Caspar, D.L., 1992. Correlations of atomic movements in lysozyme crystals. Proteins 12, 145–157. doi:10.1002/prot.340120208

Colliex, C., Cowley, J.M., Dudarev, S.L., Fink, M., Gjønnes, J., Hilderbrandt, R., Howie, A., Lynch, D.F., Peng, L.M., Ren, G., Ross, A.W., Smith, V.H., Spence, J.C.H., Steeds, J.W., Wang, J., Whelan, M.J., Zvyagin, B.B., 2006. Electron diffraction, International Tables for Crystallography. doi:10.1107/97809553602060000593

Cowtan, K., 2010. Recent developments in classical density modification. Acta Crystallogr D Biol Crystallogr 66, 470–478. doi:10.1107/S090744490903947X

DeLaBarre, B., Brunger, A.T., 2006. Considerations for the refinement of low-resolution crystal structures. Acta Crystallogr D Biol Crystallogr 62, 923–932. doi:10.1107/S0907444906012650

DeLaBarre, B., Brunger, A.T., 2003. Complete structure of p97/valosin-containing protein reveals communication between nucleotide domains. Nat. Struct. Biol. 10, 856–863. doi:10.1038/nsb972

Emsley, P., Cowtan, K., 2004. Coot: model-building tools for molecular graphics. Acta Crystallogr D Biol Crystallogr 60, 2126–2132.

Erickson, H.P., Klug, A., 1970. Fourier transform of an electron micrograph: effects of defocussing and aberrations, and implications for the use of underfocus contrast enhancement. Bericht Bunsen Gesell 74, 1129–&.

Falke, S., Tama, F., Brooks, C.L., Gogol, E.P., Fisher, M.T., 2005. The 13 angstroms structure of a chaperonin GroEL-protein substrate complex by cryo-electron microscopy. J Mol Biol 348, 219–230. doi:10.1016/j.jmb.2005.02.027

Faruqi, A.R., Henderson, R., 2007. Electronic detectors for electron microscopy. Curr Opin Struct Biol 17, 549–555. doi:10.1016/j.sbi.2007.08.014

Fernandez-Leiro, R., Scheres, S., 2016. A pipeline approach to single-particle processing in RELION. bioRxiv. doi:10.1101/078352

Fromm, S.A., Bharat, T.A.M., Jakobi, A.J., Hagen, W.J.H., Sachse, C., 2015. Seeing tobacco mosaic virus through direct electron detectors. J Struct Biol 189, 87–97. doi:10.1016/j.jsb.2014.12.002

Gabashvili, I.S., Agrawal, R.K., Spahn, C.M., Grassucci, R.A., Svergun, D.I., Frank, J., Penczek, P., 2000. Solution structure of the E. coli 70S ribosome at 11.5 A resolution. Cell 100, 537–549.

Grigorieff, N., Ceska, T.A., Downing, K.H., Baldwin, J.M., Henderson, R., 1996. Electron-crystallographic refinement of the structure of bacteriorhodopsin. J Mol Biol 259, 393–421. doi:10.1006/jmbi.1996.0328

Grosse-Kunstleve, R.W., Sauter, N.K., Moriarty, N.W., Adams, P.D., IUCr, 2002. The Computational Crystallography Toolbox: crystallographic algorithms in a reusable software framework. J Appl Crystallogr 35, 126–136. doi:10.1107/S0021889801017824

Harauz, G., van Heel, M., 1986. Exact filters for general geometry three dimensional reconstruction. OPTIK. 73, 146–156.

Henderson, R., 1992. Image contrast in high-resolution electron microscopy of biological macromolecules: TMV in ice. Ultramicroscopy 46, 1–18.

Hoffmann, N.A., Jakobi, A.J., Moreno-Morcillo, M., Glatt, S., Kosinski, J., Hagen, W.J.H., Sachse, C., Müller, C.W., 2015. Molecular structures of unbound and transcribing RNA polymerase III. Nature 528, 231–236. doi:10.1038/nature16143

Hohn, M., Tang, G., Goodyear, G., Baldwin, P.R., Huang, Z., Penczek, P.A., Yang, C., Glaeser, R.M., Adams, P.D., Ludtke, S.J., 2007. SPARX, a new environment for Cryo-EM image processing. J Struct Biol 157, 47–55. doi:10.1016/j.jsb.2006.07.003

Jacobson, R.A., Wunderlich, J.A., 1961. The crystal and molecular structure of cellobiose. Acta Crystallographica 14, 598–607. doi:10.1107/s0365110x61001893

Jiang, J., Brunger, A., 1994. Protein hydration observed by X-ray diffraction. Solvation properties of penicillopepsin and neuraminidase crystal structures. J Mol Biol 243, 100–115.

Jonic, S., 2017. Computational methods for analyzing conformational variability of macromolecular complexes from cryo-electron microscopy images. Curr Opin Struct Biol 43, 114–121. doi:10.1016/j.sbi.2016.12.011

Joseph, A.P., Malhotra, S., Burnley, T., Wood, C., Clare, D.K., Winn, M., Topf, M., 2016. Refinement of atomic models in high resolution EM reconstructions using Flex-EM and local assessment. Methods 100, 42–49. doi:10.1016/j.ymeth.2016.03.007

Kühlbrandt, W., 2014. The Resolution Revolution. Science 343, 1443–1444. doi:10.1126/science.1251652

Liao, M., Cao, E., Julius, D., Cheng, Y., 2013. Structure of the TRPV1 ion channel determined by electron cryo-microscopy. Nature 504, 107–112. doi:10.1038/nature12822

Lunin, V.Y., Afonine, P.V., Urzhumtsev, A.G., 2002. Likelihood-based refinement. I. Irremovable model errors. Acta Crystallogr., A, Found. Crystallogr. 58, 270–282.

Lyumkis, D., Brilot, A.F., Theobald, D.L., Grigorieff, N., 2013. Likelihood-based classification of cryo-EM images using FREALIGN. J Struct Biol 183, 377–388. doi:10.1016/j.jsb.2013.07.005

McMullan, G., Faruqi, A.R., Henderson, R., 2016. Direct Electron Detectors. Meth Enzymol 579, 1–17. doi:10.1016/bs.mie.2016.05.056

Merk, A., Bartesaghi, A., Banerjee, S., Falconieri, V., Rao, P., Davis, M.I., Pragani, R., Boxer, M.B., Earl, L.A., Milne, J.L.S., Subramaniam, S., 2016. Breaking Cryo-EM Resolution Barriers to Facilitate Drug Discovery. Cell 165, 1698–1707. doi:10.1016/j.cell.2016.05.040

Moriya, T., Saur, M., Stabrin, M., Merino, F., Voicu, H., Huang, Z., Penczek, P.A., Raunser, S., Gatsogiannis, C., 2017. _High-resolution Single Particle Analysis from Electron Cryo-microscopy Images Using SPHIRE. JoVE Journal of Visualized Experiments.

Morris, R.J., Blanc, E., Bricogne, G., 2004. On the interpretation and use of <|E|2>(d*) profiles. Acta Crystallogr D Biol Crystallogr 60, 227–240. doi:10.1107/S0907444903025538

Murshudov, G.N., 2016. Chapter Eleven-Refinement of Atomic Structures Against cryo-EM Maps. Meth Enzymol 277–305. doi:10.1016/bs.mie.2016.05.033

Murshudov, G.N., Vagin, A.A., Dodson, E.J., 1997. Refinement of macromolecular structures by the maximum-likelihood method. Acta Crystallogr D Biol Crystallogr 53, 240–255. doi:10.1107/S0907444996012255

Painter, J., Merritt, E.A., 2006. Optimal description of a protein structure in terms of multiple groups undergoing TLS motion. Acta Crystallogr D Biol Crystallogr 62, 439–450. doi:10.1107/S0907444906005270

Pettersen, E.F., Goddard, T.D., Huang, C.C., Couch, G.S., Greenblatt, D.M., Meng, E.C., Ferrin, T.E., 2004. UCSF Chimera‐‐a visualization system for exploratory research and analysis. J Comput Chem 25, 1605–1612. doi:10.1002/jcc.20084

Read, R.J., 1986. Improved Fourier coefficients for maps using phases from partial structures with errors. Acta Crystallogr., A, Found. Crystallogr. 42, 140–149. doi:10.1107/s0108767386099622

Rosenthal, P.B., Henderson, R., 2003. Optimal determination of particle orientation, absolute hand, and contrast loss in single-particle electron cryomicroscopy. J Mol Biol 333, 721–745.

Saxton, W.O., Baumeister, W., 1982. The correlation averaging of a regularly arranged bacterial cell envelope protein. J Microsc 127, 127–138.

Scheres, S.H.W., 2012. A Bayesian view on cryo-EM structure determination. J Mol Biol 415, 406–418. doi:10.1016/j.jmb.2011.11.010

Schmid, M.F., Sherman, M.B., Matsudaira, P., Tsuruta, H., Chiu, W., 1999. Scaling structure factor amplitudes in electron cryomicroscopy using X-Ray solution scattering. J Struct Biol 128, 51–57. doi:10.1006/jsbi. 1999.4173

Stehle, T., Harrison, S.C., 1996. Crystal structures of murine polyomavirus in complex with straight-chain and branched-chain sialyloligosaccharide receptor fragments. Structure/Folding and Design 4, 183–194.

Subramaniam, S., Kühlbrandt, W., Henderson, R., 2016. CryoEM at IUCrJ: a new era. IUCrJ 3, 3–7. doi:10.1107/S2052252515023738

Unwin, N., 2005. Refined structure of the nicotinic acetylcholine receptor at 4A resolution. J Mol Biol 346, 967–989. doi:10.1016/j.jmb.2004.12.031

van Heel, M., Keegstra, W., Schutter, W., Van Bruggen, E., 1982. Arthropod hemocyanin structures studied by image analysis. Life Chemistry Reports.

Wang, B.C., 1985. Resolution of phase ambiguity in macromolecular crystallography. Meth Enzymol 115, 90–112. doi:10.1016/0076- 6879(85)15009-3

Wang, R.Y.-R., Song, Y., Barad, B.A., Cheng, Y., Fraser, J.S., DiMaio, F., 2016. Automated structure refinement of macromolecular assemblies from cryo-EM maps using Rosetta. eLife 5, 352. doi:10.7554/eLife.17219

Wang, Z., Hryc, C.F., Bammes, B., Afonine, P.V., Jakana, J., Chen, D.-H., Liu, X., Baker, M.L., Kao, C., Ludtke, S.J., Schmid, M.F., Adams, P.D., Chiu, W., 2014. An atomic model of brome mosaic virus using direct electron detection and real-space optimization. Nat Commun 5, 4808. doi:10.1038/ncomms5808

Wheatley, R.W., Juers, D.H., Lev, B.B., Huber, R.E., Noskov, S.Y., 2015. Elucidating factors important for monovalent cation selectivity in enzymes: E. coli β-galactosidase as a model. Phys Chem Chem Phys 17, 10899–10909. doi:10.1039/c4cp04952g

Wilson, A., 1942. Determination of absolute from relative X-ray intensity data. Nature 150, 152–152. doi:10.1038/150152a0

Yonekura, K., Maki-Yonekura, S., Namba, K., 2003. Complete atomic model of the bacterial flagellar filament by electron cryomicroscopy. Nature 424, 643–650. doi:10.1038/nature01830

Zhang, K.Y., Cowtan, K., Main, P., 1997. Combining constraints for electron-density modification. Meth Enzymol 277, 53–64.

Zhao, J., Benlekbir, S., Rubinstein, J.L., 2015. Electron cryomicroscopy observation of rotational states in a eukaryotic V-ATPase. Nature 521, 241–245. doi:10.1038/nature14365

## References

[1] Bartesaghi, A. et al., Science 348, 1147–1151 (2015)

[2] Liao, M. et al., Nature 504, 107–112 (2013)

[3] Hoffmann, N. A. et al., Nature 528, 231–236 (2015)

[4] Bai, X.-C. et al., Nature 525, 212–217 (2015).

